# Characterization of the SARS-CoV-2 S Protein: Biophysical, Biochemical, Structural, and Antigenic Analysis

**DOI:** 10.1101/2020.06.14.150607

**Authors:** Natalia G. Herrera, Nicholas C. Morano, Alev Celikgil, George I. Georgiev, Ryan J. Malonis, James H. Lee, Karen Tong, Olivia Vergnolle, Aldo B. Massimi, Laura Y. Yen, Alex J. Noble, Mykhailo Kopylov, Jeffrey B. Bonanno, Sarah C. Garrett-Thomson, David B. Hayes, Robert H. Bortz, Ariel S. Wirchnianski, Catalina Florez, Ethan Laudermilch, Denise Haslwanter, J. Maximilian Fels, M. Eugenia Dieterle, Rohit K. Jangra, Jason Barnhill, Amanda Mengotto, Duncan Kimmel, Johanna P. Daily, Liise-anne Pirofski, Kartik Chandran, Michael Brenowitz, Scott J. Garforth, Edward T. Eng, Jonathan R. Lai, Steven C. Almo

## Abstract

Coronavirus disease 2019 (**COVID-19**) is a global health crisis caused by the novel severe acute respiratory syndrome coronavirus 2 (**SARS-CoV-2**), and there is a critical need to produce large quantities of high-quality SARS-CoV-2 Spike (**S**) protein for use in both clinical and basic science settings. To address this need, we have evaluated the expression and purification of two previously reported S protein constructs in Expi293F^™^ and ExpiCHO-S^™^ cells, two different cell lines selected for increased expression of secreted glycoproteins. We show that ExpiCHO-S^™^ cells produce enhanced yields of both SARS-CoV-2 S proteins. Biochemical, biophysical, and structural (**cryo-EM**) characterization of the SARS-CoV-2 S proteins produced in both cell lines demonstrate that the reported purification strategy yields high quality S protein (non-aggregated, uniform material with appropriate biochemical and biophysical properties). Importantly, we show that multiple preparations of these two recombinant S proteins from either cell line exhibit identical behavior in two different serology assays. We also evaluate the specificity of S protein-mediated host cell binding by examining interactions with proposed binding partners in the human secretome. In addition, the antigenicity of these proteins is demonstrated by standard ELISAs, and in a flexible protein microarray format. Collectively, we establish an array of metrics for ensuring the production of high-quality S protein to support clinical, biological, biochemical, structural and mechanistic studies to combat the global pandemic caused by SARS-CoV-2.

---

Most human coronavirus infections are associated with mild symptoms, but in the last two decades, three beta coronaviruses, SARS-CoV, MERS, and SARS-CoV-2, have emerged that are able to infect humans and cause severe disease [1, 2]. The current pandemic of coronavirus disease 19 (**COVID-19**) is caused by severe acute respiratory syndrome coronavirus 2 (**SARS-CoV-2**) [3], an enveloped virus from the *Coronaviridae* family with a single positively stranded RNA genome [3]. This RNA virus, which likely originated in bats, has several structural components, including Spike (**S**), Envelope (**E**), Membrane (**M**), and Nucleocapsid (**N**) proteins [2].

The S protein is a class I viral fusion protein, which consists of two subunits (**S1** and **S2**) and forms a trimer on the viral membrane [4]. The S1 subunit contains the receptor binding domain (**RBD**) which is responsible for host cell receptor binding, while the S2 subunit facilitates membrane fusion between the viral and host cell membranes [4-7]. Host cell proteases are essential for activating the S protein for *Coronaviridae* cellular entry [8]. The S protein in many *Coronaviridae*, including SARS-CoV-2, is cleaved between the S1 and S2 subunits (S1/S2 cleavage site) and at an additional site present in S2 (S2’ cleavage site) [8-10]. Similar to SARS-CoV, the SARS-CoV-2 trimeric S glycoprotein mediates viral entry into the cell by utilizing angiotensin converting enzyme 2 (**ACE2**) as a human cell surface entry receptor [8]. Processing of both the SARS-CoV and SARS-CoV-2 S protein is dependent on the endosomal cysteine proteases cathepsin B and cathepsin L, and the serine protease ™PRSS2[8]. In many coronaviruses, these events lead to conformational rearrangements in S2, which ultimately result in fusion of the host and viral cell membranes, and delivery of the viral genome into the newly infected cell [11].

Due to the global COVID-19 pandemic, the SARS-CoV-2 S ectodomain protein has become an important target for clinical, biological and structural investigations, and future studies will require the efficient and streamlined production of this protein. Clinically, the production of large amounts of S ectodomain protein enables testing of individuals for SARS-CoV-2 seropositivity. Serological testing is important for determining individuals who have been exposed to the virus, and the resulting antibody titer can facilitate identification of potential donors of convalescent plasma[12]. Additionally, the S protein ectodomain could be used to identify therapeutic monoclonal antibodies (**mAbs**) through single B cell cloning from convalescent patients. Furthermore, the development of small molecules and protein therapeutics designed to inhibit viral infection by targeting the S protein need to be tested and biochemically characterized. Biologically, the complete mechanisms of viral host cell fusion and replication remain to be completely understood. Structural studies will continue to support these ongoing clinical, therapeutic and biological investigations.

To provide support for these investigations, we examined the expression and purification of two recently reported recombinant versions of the S protein (here termed **OptSpike1** and **OptSpike2**, see **Fig. 1A**). OptSpike1 was initially reported by McLellan and coworkers [7], and used to determine the cryo-EM structure of the Spike protein in the prefusion conformation and has been utilized as an antigen for clinical ELISAs at Montefiore Medical Center (Bronx, NY) [**Bortz et al**., **manuscript in preparation**]. OptSpike2 was described by Krammer and coworkers, and has been successfully used to conduct serum ELISAs to test patients for the presence of anti-S antibodies at Mount Sinai Hospital (New York, NY) [13]. The successful use of both of these recombinant forms of the S protein in clinical (OptSpike1 and OptSpike2) and structural (OptSpike1) applications have made them attractive targets for future COVID-19 studies. Both constructs are cloned into the mammalian expression vector pCAGGS, and include either the majority of the ectodomain (**OptSpike1: AAs 1-1208**) or the full-length ectodomain (**OptSpike2: 1-1213**) of the SARS-CoV-2 S protein (based on the Wuhan-Hu-1 sequence) [14]. Both constructs include the K986P and K987P stabilizing mutations and use a T4 Foldon motif (**T4**) to enhance trimerization [15, 16]. Both constructs lack the furin cleavage site: OptSpike1 contains the mutation RRAR:GSAS, while OptSpike2 contains the mutation RRAR:A [17]. OptSpike1 contains the T4-HRV3C protease cleavage sequence-8x HisTag and a TwinStrepTag at the C-terminus, while OptSpike2 contains a Thrombin cleavage sequence-T4-6x HisTag at the C-terminus.

**Figure 1:**
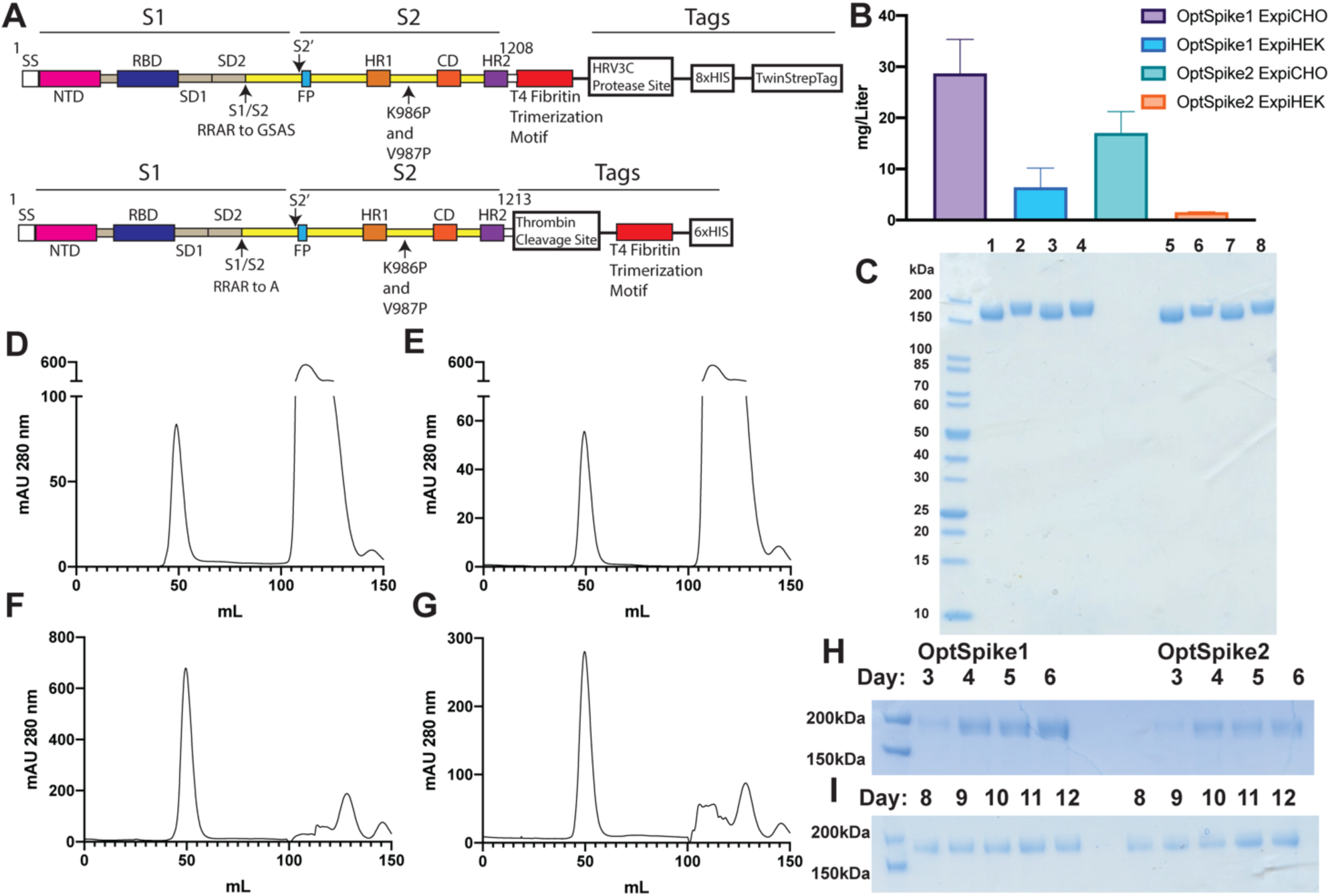
Expression and Purification of the SARS-CoV-2 S Protein. **A)** Schematic showing characteristic of OptSpike1 (upper) and OptSpike2 (lower) **B)** Average yields of OptSpike1 and OptSpike2 produced in either Expi293F^™^ or ExpiCHO-S^™^ cells harvested on either day 6 (HEK) or day 12 (CHO), and purified using Nickel affinity chromatography and size exclusion chromatography **C)** SDS-PAGE showing apparent molecular mass and purity of OptSpike1 and OptSpike2 purifications. Lanes 1-4: 3 µg OptSpike1-CHO, OptSpike1-HEK, OptSpike2-CHO, OptSpike2-HEK in the presence of 100 μM DTT, Lanes 5-8: Same as 1-4 without DTT **D-E)** Representative HiLoad^™^ 16/600 Superdex^™^ 200 purification of **D)** OptSpike1-HEK **E)** OptSpike2-HEK **F)** OptSpike1-CHO **G)** OptSpike2-CHO after nickel affinity purification **H)** Yield from 1 mL crude nickel purification of OptSpike1 or OptSpike2 on indicated day post transfection in Expi293F cells **I)** Yield from 1 mL crude nickel purification of OptSpike1 or OptSpike2 on indicated day post transfection in ExpiCHO-S cells.

Here, we examine the expression of OptSpike1 and OptSpike2 in Expi293F^™^ and ExpiCHO-S^™^ expression systems [18], and provide detailed protein production protocols (including an SOP in **Appendix 1**). We then evaluate the biochemical and biophysical properties of these proteins to define standards of protein quality and evaluate the recognition specificity of the S protein within the human secretome. Furthermore, we demonstrate the functional reproducibility of recombinant S protein in serum ELISAs and develop a multi antigen COVID-19 protein microarray that can simultaneously test for antibodies against multiple antigens, including the full-length S protein, the receptor binding domain (**RBD**) of the S protein, and the full-length N protein. Finally, we determine the 3.22 Å cryo-EM structure of OptSpike1 produced in ExpiCHO-S^™^ cells, and the 3.44 Å cryo-EM structure of OptSpike1 produced in Expi293F^™^ cells, which are nearly indistinguishable and in agreement with previously described structures [7, 19].

This work provides a comprehensive strategy, with a wide range of guiding standards and metrics, for production of large quantities of high-quality recombinant S protein for use in future clinical, diagnostic, biological, biochemical, structural and mechanistic studies that will be needed to combat the global pandemic caused by SARS-CoV-2.

## Materials and Methods

### Recombinant spike protein transfection

#### An SOP for the production of high-quality recombinant S antigen is provided in Appendix 1

Both variants of the recombinant SARS-CoV-2 S protein (OptSpike1 and OptSpike2) were expressed in Expi293F^™^ and ExpiCHO-S^™^ cells according to manufacturer’s instructions (ThermoFisher Scientific). For Expi293F^™^ (ThermoFisher Cat# A14527) purifications, cells were grown in sterile TC flasks, vented, with a baffled bottom (example: FisherScientific Cat # BBV12-5) in a Climo Shaker ISF4-X at 110 RPM (orbital diameter of 50 mm) at 37° C and 8% CO2. When cells reached a density between 3.5 and 5×10^6^ cells / mL, they were counted, and appropriately diluted into a shaker flask. For example, for a transfection volume of 30 mL, 7.5 × 10^7^ cells were brought to a total volume of 25.5 mL in a 125 mL shaker flask. Cells were then transfected by diluting 30 μg of plasmid DNA in Opti-MEM(tm) I Reduced Serum Medium (Cat # 31985-062) to a total volume of 1.5 mL in a 15 mL conical tube, and then mixing briefly by inverting the tube. 80 μL of ExpiFectamine(tm) 293 Reagent was then diluted in Opti-MEM(tm) I medium to a total volume of 1.5 mL, in a separate 15 mL conical tube. Both reactions were incubated for 5 minutes. The transfection reagent mixture was then carefully pipetted into the DNA mixture, and mixed by inverting the 15 mL conical tube 3 times. The combined DNA and transfection reagent mixture was incubated at room temperature for 25 minutes, after which the complexes were added to the cell culture in a drop wise fashion, while swirling the cell culture flask to ensure uniform distribution of the DNA complexes. 16 hours post-transfection, 150 μL of ExpiFectamine(tm) 293 Transfection Enhancer 1 and 1.5 mL of ExpiFectamine(tm) 293 Transfection Enhancer 2 were added to the transfected cells, bringing the total volume of the transfection to 30 mL. Expi293F^™^ transfections were harvested on day 6 post transfection unless otherwise specified. This protocol can be scaled linearly for larger Expi293F^™^ cultures, according to the manufacturer’s protocol, for both OptSpike1 and OptSpike2.

For ExpiCHO-S^™^ transfections, we obtained the ExpiCHO-S(tm) (ThermoFisher Cat # A29127) cells and ExpiCHO Expression System kit (Cat # A29133) from ThermoFisher Scientific. ExpiCHO-S(tm) cells were grown in sterile, Erlenmeyer-shaped flasks with plain bottom with vented screw caps (appropriate flasks for cell culture volume should be used see for example: Fisherbrand PBV125) in a Climo Shaker ISF4-X at 110 RPM (orbital diameter of 50 mm) at 37° C and 8% CO2. ExpiCHO-S(tm) cells were passaged and split every 2-3 days when cell densities reached 0.3×10^6^-6×10^6^ viable cells per mL. ExpiCHO-S(tm) cells are allowed to reach a density of 7-10×10^6^ cells per mL on the day of transfection, with a viability of at least 95%. ExpiCHO-S cells are diluted to a final density of 6×10^6^ cells per mL in a plain bottom Erlenmeyer flask. For spike protein transfections, the manufacturers ExpiCHO expression system manual was followed closely. For example, for a 25mL transfection, 25 μg of filtered DNA was diluted into 1mL of OptiPRO^™^ SFM (Cat # 12309019) and 80 μL of ExpiFectamine^™^ CHO reagent was added to 920 μL of OptiPRO^™^ SFM. Dilutions were mixed by inversion and then diluted DNA was mixed with diluted ExpiFectamine^™^ CHO reagent and mixed by inversion. ExpiFectamine^™^ CHO and DNA complexes were incubated at room temperature for 5 minutes and slowly transferred to a 125mL plain bottom flask containing 25mL of ExpiCHO-S cells at a cell density of 6×10^6^ cells per mL. For all spike transfections, we followed the max titer protocol (unless otherwise stated), thus on day 1 post transfection we added 150 μL of ExpiCHO-S^™^ Enhancer and 4mL of ExpiCHO-S^™^ Feed. On day 1 post transfection, the cells were shifted to another Climo Shaker ISF4-X at 110 RPM (orbital diameter of 50 mm) with temperature set at 32° C and 5% CO2. On day 5 post transfection, another 4mL of ExpiCHO-S^™^ Feed is added and the cells are set back at 32° C. ExpiCHO-S^™^ transfections were harvested on day 12 post transfection unless otherwise specified.

### S Protein Purification

For purifications of the SARS-CoV-2 S Protein (OptSpike1 and OptSpike2), on the indicated day post transfection (optimally day 6 for Expi293F(tm) and day 12 for ExpiCHO-S(tm)), the media was harvested by centrifuging cells at 500g for 10 minutes, removing the supernatant, and centrifuging the supernatant at 2000g for 10 minutes. For ExpiCHO-S(tm), the supernatant was clarified by filtering through a rapid flow filter from ThermoFisher (example: 524-0020). The supernatant was then adjusted to 50 mM ArgCl using a 2 M stock solution of ArgCl at pH 6.5, to increase stability of the protein during purification. OptSpike1 and OptSpike2 were subsequently purified by His60 Ni^2+^ superflow resin (Takara Cat: 635664) using a batch binding method: .6 mL of resin were added for every 10 mL of supernatant, and supernatants were incubated with resin for 2 hours at 4° C with shaking or rotating. Batch adsorption was followed by gravity flow over a column. The Ni^2+^His60 resin was washed with 3 column volumes of wash buffer (50 mM TRIS pH 8.0, 100 mM ArgCl, 5 mM imidazole, 150 mM NaCl, 10% Glycerol) and the bound protein eluted with 5 mL of the same buffer containing 500 mM imidazole. Nickel eluates were concentrated by centrifugation in 100K concentrator (ThermoFisher Cat # 88533) by spinning at 1100 g for intervals of 8 minutes (if necessary) and further purified by gel filtration on a HiLoad^™^ 16/600 Superdex^™^ 200 column (GE) equilibrated with 50 mM TRIS, 100 mM ArgCl, 150 mM NaCl, 10% Glycerol, pH 8.0. For use in cryo-EM experiments, nickel eluates were further purified by gel filtration on a HiLoad^™^ 16/600 Superdex^™^ 200 (GE) equilibrated with 50 TRIS, 150 mM NaCl, pH 8.0. For protein used in serum ELISA or protein microarray experiments, 30 ml of Nickel eluates were dialyzed against 5 L of 50 mM TRIS, 150 mM NaCl, pH 8.0 at 4 °C for 16 hours. Generally, the SEC step did not affect the serum ELISA experiments, but was important for all other experiments. Protein concentration was determined using an extinction coefficient (1468500 M^-1^cm^-1^) estimated from amino acid sequence by Expasy online ProtParam tool[20].

### SARS-CoV2 RBD Expression and Purification

The pCAGGs SARS-CoV2 RBD plasmid (provided by Florian Krammer) was used for recombinant RBD expression as previously described. FreeStyle 293F cells (ThermoFisher, R79007) were transiently transfected with a mixture of plasmid DNA diluted in PBS (0.67 μg total plasmid DNA per ml of culture) and Polyethylenimine (PEI) (Polysciences, Inc., 23966) at a DNA-to-PEI ratio of 1:3. At six days post-transfection, cultures were harvested by centrifugation at 4,000 x g for 20 min, and supernatant was incubated with Ni-NTA resin (Goldbio) for 2 hours at 4° C with gentle stirring. Resin was collected in columns by gravity flow, washed with 16 column volumes of wash buffer (50 mM Tris HCl pH 8.0, 250 mM NaCl, 20 mM Imidazole) and eluted in 12mL elution buffer (50 mM Tris HCl pH 8.0, 250 mM NaCl, 250 mM Imidazole). Eluates were concentrated and exchanged into storage buffer (50 mM Tris HCl pH 8.0, 250 mM NaCl) using an Amicon centrifugal units (EMD Millipore). Protein concentration was determined using an extinction coefficient (33350 M^-1^cm^-1^), estimated from amino acid sequence by Expasy online ProtParam, and was further analyzed by SDS-PAGE.

### Recombinant expression and purification of SARS-COV-2 N-Protein

The nucleocapsid sequence was PCR amplified from a diagnostic test positive control plasmid obtained from IDTDNA, and InFusion cloned into a derivative of the NYSGRC pSGC-HIS vector. 50 ng of plasmid was used to transform 20 μl of the BL21 DE3 strain of *E. coli*. Cultures were then grown overnight (16 hours) in LB at 37°C and used to inoculate either LB media the next day (1:100x dilution of overnight culture). Inoculated cultures were grown at 37°C until they reached OD_600_ 0.7, at which point they were induced using 500 μM IPTG. Upon induction of LB media, temperature of the cultures was immediately lowered to 25°C for 16 hours.

To harvest protein, cells were lysed by sonic disruption using a 550 sonic dismembrator from Fisher Scientific. Every 5 g of were resuspended in 30 mL of lysis buffer consisting of 50 mM HEPES, 250 mM KCl, 10% glycerol, 10 mM BME, 0.1% Igepal® CA-630 (Sigma Aldrich), pH 7.5 and ½ protease inhibitor tablets (Roche). After lysis, samples were cleared by centrifugation at 20,000 rpm. The resulting supernatant was purified on an AKTA FPLC (GE Biosciences). Supernatants were loaded onto fast flow HisTrap columns and washed with 20 column volumes of lysis buffer and eluted with 2 column volumes of Buffer B (Buffer A + 500 mM imidazole, pH 7.5). The resulting eluent with high OD_280_ absorbance was collected and loaded onto a HiPrep 16/60 S-200 size exclusion column equilibrated with 50 mM HEPES, 250 mM KCl, 10% glycerol, 5 mM DTT, pH 7.5. Protein concentration of fractions were approximated using an extinction coefficient of 43890 M^-1^cm^-1^, and molecular mass of 45.62570 kDa estimated from amino acid sequence by Expasy online ProtParam tool[20].

### Analytical Size Exclusion Chromatography

After nickel elutes were concentrated and purified by gel filtration on a HiLoad^™^ 16/600 Superdex^™^ 200 column and concentrated, protein aggregation state was assessed by analytical gel filtration on a Superose^™^ 6 Increase 10/300 GL column. The void for this column runs at 8.5mL. Aggregation state was monitored over time, and after freeze thaw cycles on this column.

### Molecular Mass Determination using Multi Angle Light Scattering (MALS)

30 μL of either OptSpike1, Optspike2, or Nucleocapsid was run over a Yarra(tm) 3 µm SEC-4000 LC Column using an Agilent Technologies 1260 Infinity instrument, equipped with auto injector. 10 µl samples were injected onto the column in 50 mM Tris, 150 mM NaCl, 100 mM ArgCl, 10% glycerol, pH 8.0 at a flow rate of 0.25ml/min. MALS analysis was performed using miniDAWN Treos MALS detector (Wyatt) and Optilab T-rEX and analyzed using the associated Astra software. Baselines were determined automatically; peaks were manually delineated.

### OptSpike1 Protein Melting Curve

5000x stock of SYPRO^™^ dye was diluted to 200x in Tris-HCl (pH 7.5), 100 mM KCl buffer, the same buffer as OptSpike1 and OptSpike2 were diluted in for the assay. Stock protein was serially diluted to 1.75 μM; 45 μl of 1.75 μM protein was aliquoted into 1.5 mL Eppendorf tubes, and 5 μl of 200x dye was added to each tube. The final concentration of protein was 1.58 μM and 20x SYPRO^™^ Orange dye. The samples were split into three technical triplicates (one in each well of a 384 well plate), and the reactions were placed into 7900 HT Fast Real Time PCR System. The PCR machine was programmed to monitor the fluorescence of the dye over a temperature gradient spanning 25°C to 99°C. The protocol was designed to hold the samples at 25°C for two minutes, and then ramp the temperature Background fluorescence was measured using 20x dye and all measurements were taken in technical triplicate.

Data was exported in Excel, and then analyzed and graphed by GraphPad Prism. To discern the melting temperature (**™**) of the protein, two methods were used as outlined in: 1) fitting the Boltzmann equation to the non-linear raw data and 2) detection of local maximum values in the 1^st^ derivative of melting curve. The curve had two apparent melting transitions, one between 42-55°C and 55-70°C. The curves in these two temperature ranges were fitted with Boltzmann equations (listed below). The function fit to data in each of the temperature ranges contained an approximate value for each ™. In order to determine the local maxima values the 1^st^ and 2^nd^ derivatives were graphed (the 2^nd^ derivative should equal 0 at the local maxima of the 1^st^ derivative, which can be visually verified by the graph of the 1^st^ derivative). Both methods produced comparable melting temperature values. OptSpike1 ™_1_ = 49.04±0.1699°C and OptSpike2 ™_1_ = 49.36±0.1652°C; OptSpike1 ™_2_ = 63.04 ± 0.1659°C and OptSpike2 ™_2_ = 63.31 ± 0.09236°C. The melting temperature can also be approximated by determining local maxima of the 1^st^ derivative of Fluorescence vs temperature curve. There wasn’t a significant difference in either ™_1_ and ™_2_ between the OptSpike1 and OptSpike2 constructs.

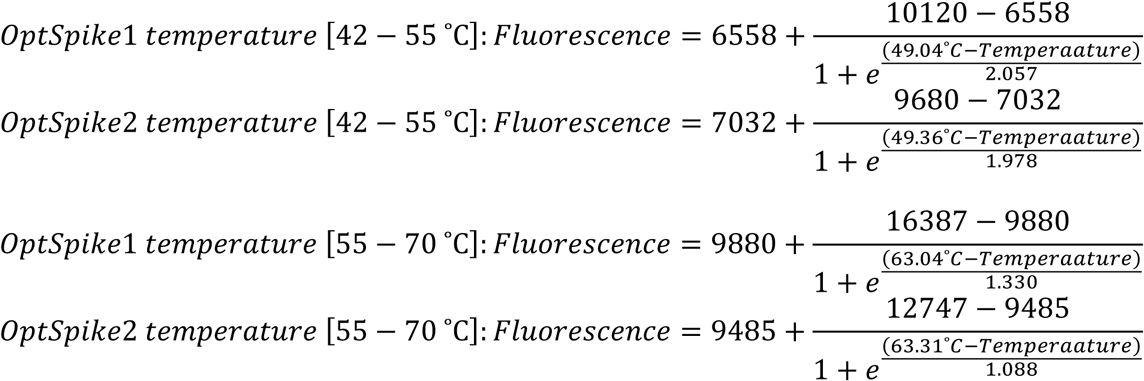

### Analytical Ultracentrifugation of OptSpike1

Analytical ultracentrifugation (**AUC**) studies were conducted with OptSpike1 in 250mM NaCl 50mM Tris pH 8.0 in a Beckman XL-AUC using the absorption optics to scan cells assembled with a double sector charcoal EPON centerpiece and sapphire glass windows inserted into a AN-60 Ti rotor. Centrifuge runs were conducted at 20 °C and 30,000 rpm. The first run was scanned at 230 nm. A second run was scanned at 280 nm. This protocol allowed protein concentrations ranging from 136 nM to 4.8 μM to be analyzed. A minimum of 50 scans were acquired for each sample. An overview of the sedimentation behavior of each sample was obtained by time-derivative g(s*) analysis conducted using either the program SedAnal [21, 22] or the program DCDT+ by John Philo v2.5.1[23, 24]. Component analysis was conducted using the program Sedfit [25, 26] to deconvolute the species present in a solution. The program Prism v8 was used to generate the AUC figures shown in the text and supplement.

### Convalescent Serum and Plasma Samples

Serum and plasma samples were collected from healthy adult volunteers residing in Westchester County, NY who had recovered from COVID-19 in April 2020. Patients had reported a positive nasopharyngeal swab by PCR for SARS-CoV-2 during illness and had been asymptomatic for at least 14 days prior to sample collection. After obtaining informed consent, serum was obtained by venipuncture (BD Vacutainer, serum), centrifuged, aliquoted and stored at -80°C prior to use. The sera were heat-inactivated at 56°C for 30 minutes and stored at 4°C prior to analysis. Protocol approval was obtained by the Institutional Review Board (protocol IRB# 2016-6137) of the Albert Einstein College of Medicine.

### Protein Microarray Production and Processing

The COVID-19 protein array included OptSpike1, OptSpike2, receptor binding domain (RBD) of Spike protein and nucleocapsid proteins, along with positive (human IgG) and negative (human acetylcholinesterase, PBS or printing buffer) controls. Arrays were generated by printing purified proteins onto aminosilane coated slides with printing buffer containing an amine-to-amine homobifunctional crosslinker (bissulfosuccinimidyl suberate, BS3, Thermo Scientific Pierce(tm) Cat # A39266) and glycerol. BS3 was used for covalent immobilization of the proteins to the slide, and glycerol to keep them hydrated at all times. Protein concentrations were adjusted to 25, 50, 100 and 200pg per spot (two-fold dilutions between 25 and 200pg per spot). The array layout was designed with sixteen identical subarrays. Each sample was spotted in duplicate with a total drop deposition volume of 800pL (8 drops of 100 pL) per spot using Marathon Argus piezoelectric printer (Arrayjet, Edinburgh, UK) at 50-55% humidity. The slides were incubated in a humidity controlled (50%) enclosure for 45 minutes and blocked by incubation with Superblock(tm) blocking buffer (Thermo Fisher Scientific, Rockford, IL) for 45 minutes at room temperature with shaking at 60rpm.

After blocking, a multi-well chamber (ProPlate® Multi-Well Chambers, Grace Bio-labs, Bend, OR) was used to screen multiple serum samples with subarrays on the same slide. As negative control for screening, we used three serum samples (1, 4 and 8) which were received from the New York Blood Center which were collected before the 2019 SARS-CoV-2 outbreak and stored as aliquots at -80 °C. COVID-19 convalescence sera samples (2, 3, 5, 6, 7) were previously validated to be negative with RT-PCR for SARS-CoV-2 and positive ELISA for antibodies against SARS-CoV-2.

Each subarray was challenged with 250 µl of 1:100 patient sera diluted in 5% (w/v) milk, PBS, 0.2% (v/v) Tween-20 (5% milk-PBST) overnight at 4 °C with gentle shaking on a plate shaker. The arrays were rinsed with 5% milk-PBST briefly and incubated for 1 hour at room temperature with a fluorescently labeled secondary antibody (Alexa Fluor 647 labeled goat anti-human IgG (H+L) cross-adsorbed secondary antibody, Thermo Fisher Scientific, Cat # A21445) diluted 1:150 in 5% milk-PBST. After washing three times with PBST, the slides were rinsed with water and dried in falcon tubes by centrifugation at 1000xg for 3 minutes. The slides were scanned with a GenePix® 4400A microarray scanner (Molecular Devices), and data were analyzed with GenePix® Pro 7 software. Mean fluorescence intensities (**MFI**) of every spot was quantified after background correction was performed for each serum sample by subtracting the MFI of huAche, which was printed as negative control within the subarray. Duplicate spot measurements were averaged. The results were normalized relative to corresponding signal of hIgG1 and multiplied by a hundred for presentation as percentage.

### Recombinant Protein ELISAs

Recombinant SARS-CoV-2 Spike protein was coated onto high-binding 96-well plates (Corning, 3690) at 2 μg/mL, overnight at 4 °C. Wash steps were done using 1xPBS containing 0.05% Tween-20 (Sigma, P1379) and all incubations were done at 25 °C. Plates were blocked with 3% nonfat dry milk (BioRad, P1379) in 1xPBS for 1 hour. Serum from COVID-19 convalescent patients was serially diluted in 1XPBS, containing 1% nonfat dry milk and 0.1% Tween-20, then added to plates and incubated for 2 hours. Following plate washing, serum reactivity to the Spike protein was measured using an HRP-conjugated goat anti-human IgG (H+L) (Invitrogen, 31410) at a 1:3000 dilution for 1 hour. After a final wash, Ultra-™B substrate (ThermoFisher, 34029) was added to the plates and incubated for 5 minutes, followed by quenching of the reaction by addition of an equal volume of 0.5M H_2_SO_4_. Absorption was measured at OD_450_ using a Synergy4 plate reader (Biotek), and data was analyzed using GraphPad Prism7.0 to calculate IC50 values. **PNGase F Digestion of Spike Protein** For PNGase F digestion, purified Spike protein expressed from Expi293F^™^ or ExpiCHO-S^™^ cells was digested under denaturing reaction conditions. 5 μg of protein, 1 μL of Glycoprotein denaturing buffer (NEB Cat # B0701S, 10X) and 4 μL water were mixed to a total of 10 μL. The spike glycoproteins were denatured at 100° C for 10 minutes, and the denatured proteins incubated on ice for 5 minutes and centrifuged for 10 seconds at max speed on a microcentrifuge. Next, 2 μL of GlycoBuffer 2 (NEB Cat # B0701S, 10X), 2 μL of 10% NP-40 (NEB Cat # B0701S) and 6 μL of water were mixed with the 10 μL of denatured spike glycoprotein. Finally, 1 μL of PNGase F (NEB Cat # P0704S) was added to the reaction and the digestion allowed to proceed at 37° C for 1 hour. Glycoproteins were analyzed by SDS-PAGE to observe loss of N-linked glycans compared to undigested glycoproteins.

### HEK293F Transient Transfections and Culture

HEK 293 suspension cells were cultured in HEK Freestyle Media (Invitrogen) at 37 C in a humidified shaking platform incubator (Kuhner) with 5% CO2. For transfection, cells were pelleted at 500xg and resuspended in fresh media. For small-scale (1mL cells at 1×106/mL) transient transfections performed in 24-well non-treated tissue culture plates, 2 μg Polyethylenimine (**PEI**) was added to 0.5 μg diluted plasmid DNA in a final volume of 100 μL. For small-scale (.25 mL cells at 1 x10^6^/mL) transient transfections performed in 48-well non-treated tissue culture plates, .5 μg Polyethylenimine (**PEI**) was added to 0.125 μg diluted plasmid DNA in a final volume of 25 μL. DNA-PEI complexes were incubated at room temperature for 10 minutes, and then added directly to cells in 48-well plates.

### Flow Cytometry Titration Experiment

Flow cytometry titration assays were performed with the OptSpike1 and OptSpike2 proteins described above. HEK293F suspension cells were transfected with human ACE2, mouse ACE2, human CD26, or GFP. Both ACE2 variants were expressed with GFP tags on their cytosolic C term, while CD26 was expressed in an IRES vector expressing GFP. Two days post transfection, cells were counted and diluted to 1×10^6 cells/mL in 1x PBS, .2% BSA. OptSpike1 and OptSpike2 proteins were serially diluted to a range of concentrations from 1 nM - 10 μM. Subsequently, 10 μL of diluted Spike protein was added to 90 μL diluted cells in wells of a 96-well plate (90,000 cells per well). Binding was performed at room temperature for 1 hour with shaking at 900 rpm, after which the cells were washed 2x with 1x PBS .2% BSA by centrifugation. Cells were then incubated with a PE-labeled anti-6x His tag antibody (Abcam Cat # ab72467), to detect Spike protein binding. Antibody binding was performed for 30 minutes at room temperature with shaking, after which the cells were washed 2x with 1x PBS, .2% BSA by centrifugation. Cells were analyzed by Flow Cytometry/Spectral Analysis on a SONY Spectral Analyzer. Gated live cells were sub-gated for GFP, and GFP positive cells sub-gated for PE positive events. Data points represent the average of three independent experiments fit to the single site binding equation: Y=Bmax*X^h^/(Kd^h^ + X^h^).

### Generation of a Human Plasma Membrane Protein Library for Mammalian Expression

The human genome was analyzed for all transcripts that contain at least one transmembrane domain (™HMM Server 2.0) and a predicted secretion signal peptide sequence [27, 28]. The resulting list of 14,028 transcripts was manually purged of mitochondrial, nuclear and ER membrane proteins, resulting in a final target set of 9,065 potential human plasma membrane transcripts representing a set of 4,860 genes. All of the full-length cDNAs available from GeneCopoeia^™^ were purchased as covalent C-terminal GFP fusions in a CMV mammalian expression plasmid (3926 Total). To enhance library coverage of proteins missing from the GeneCopoeia library, a separate set of synthetic transcripts was ordered from Gen9(tm) (1282 Total). This set was selected by identifying the largest transcript for each of the missing proteins with the limitation that they be under 4kb in length (the limit for high-throughput synthesis). Each full-length cDNA was codon optimized, synthesized and sub-cloned into the Clonetech pEGFP N1 vector as C-terminal GFP fusions. Working libraries of Ig superfamily proteins (IGSF), TNF receptor superfamily proteins (TNFRSF), G-Protein coupled receptor proteins (GPCR), Integrins and chemokines were identified for each subfamily based on lists generated using the HUGO Gene Nomenclature Committee (HGNC) gene family resource[29]. Each set was re-arrayed manually from glycerol stocks into liquid 2xYT media in 96-well deep-well blocks. Overnight cultures were mini-prepped in 96-well plates (Macherey Nagel kits) for use in downstream high-throughput mammalian cell transfections.

### High-throughput Screening of Human Plasma Membrane Protein Library

The human plasma membrane protein library was transfected into HEK293F cells in 48-well format. Two days post-transfection, cells were diluted to 1 x10^6^cells/mL in .2% BSA. Binding reactions were setup in 96-well V-bottom plates by incubating 100 μL cells with 200 nM OptSpike1; each plate also contained hACE2-GFP expressing cells as an internal positive control. After 45 minutes, cells were pelleted and washed twice, and then incubated with an anti-HIS PE antibody, washed twice, and analyzed on a SONY Spectral Analyzer. The percent bound was calculated as the number of double-positive events (GFP and mCherry) divided by the total number of GFP positive cells. Expression of hACE2, mACE2, CD147, CD26, Siglec9, Siglec10, Ceacam1, and Ceacam5 were confirmed by antibody staining of transfected HEK293F cells. Binding was conducted as described for S protein binding. Antibodies used: hACE2 and mACE2 (RND Cat # AF933-SP), CD147 (Biolegend Clone HIM6), CD26 (Biolegend Clone BA5b), Siglec9 (Biolegend Clone K8), Siglec10 (Biolegend Clone FG6), Ceacam1 and Ceacam5 (Biolegend Clone ASL-32).

### CryoEM

#### Grid Preparation

3µl of protein (1mg/mL in 50 mM TRIS, 250 mM NaCl, pH 8.0) was applied to plasma-cleaned C-flat 1.2/1.3 400 mesh Cu holey carbon grids (Protochips, Raleigh, NC) or 1.2/1.3 300 mesh UltrAuFoil gold holey gold grids (Quantifoil Micro Tools GmbH, Großlöbichau, Germany), blotted for 2.5 s after a 30 s wait time, and then plunge frozen in liquid ethane, cooled by liquid nitrogen, using the EM GP2 (Leica Microsystems, Inc, Buffalo Grove, IL) or Vitrobot Mark IV (Thermo Fisher Scientific, Hillsboro, Oregon).

#### Microscopy

Thermo-Fisher Titan Krios operated at 300 kV, Gatan GIF-Bioquantum with a 20 eV slit and K3 camera were used with a 100µm C2 aperture, 100 µm objective aperture and calibrated pixel size of 1.058Å.

#### Imaging

Movies were collected in counting mode using Leginon [30] at a dose rate of 26.6 e-/Å2/s with a total exposure time of 2.5 seconds, for an accumulated dose of 66.5 e-/Å2. Intermediate frames were recorded every 50 ms for a total of 50 frames per micrograph. Defocus values range from approximately 0.8 – 2.5 µm.

#### Image Processing

Movies recorded on the K3 were aligned using Appion[31] and MotionCor2[32], and CTF estimated and 2D classified in cryoSPARC v2.14.2[33]. Particle picking was done with TOPAZ[34] as implemented in cryoSPARC. A Topaz picking model was trained using frame-summed micrographs of 4,000 particles manually curated from 100 micrographs initially picked from 7,500 blob picks in cryoSPARC. For the final reconstruction, particles were selected and subjected to 3D refinement in cryoSPARC with a final box size of 384×384pixels. The CHO expressed dataset (n20apr21a) processing used 1,131 micrographs and 75,582 particles that was curated to 54,395 particles. The HEK expressed dataset (n20apr22) processing used 1,694 micrographs and 99,154 particles that was curated to 54,066 particles.

## RESULTS

### Enhanced expression and purification of SARS-CoV-2 S proteins

To determine conditions for the enhanced expression and purification of the SARS-CoV-2 S protein, we tested the expression of two recently reported recombinant variants (**OptSpike1** and **OptSpike2, Fig. 1A**) in both the Expi293F^™^ and ExpiCHO-S^™^ cells [18]. The ExpiCHO-S^™^ expression system reproducibly afforded the highest yield per liter (∼28 mg/L) of transfected cells for both versions of the S protein (**Fig. 1B**). OptSpike1 and OptSpike2 were readily produced at a high level of purity, via transient transfection of both Expi293F^™^ and ExpiCHO-S^™^ cells, followed by purification with Nickel affinity and size exclusion chromatography (**SEC**) (**Fig. 1C**). SEC revealed that both S proteins ran at the appropriate size on a HiLoad^™^ 16/600 Superdex^™^ 200 gel filtration column (**Fig. 1D-G**), without the presence of significant aggregates. To identify optimal culture growth times, a time course was performed for both constructs in both expression systems, and the optimal day for harvest was found to be day 6 in Expi293F^™^ cells and day 12 in ExpiCHO-S^™^ cells (**Fig. 1H-I**). Additionally, the ExpiCHO-S^™^ standard titer and high titer protocols (**Fig. S1H**) were compared. Collectively, these data reveal that the ExpiCHO-S^™^ expression system is more efficient than the Expi293F^™^ expression system for the production of recombinant Spike protein. To provide further information about the stability, quality, and aggregation state of OptSpike1 and OptSpike2, their biophysical properties were characterized by a number of approaches.

### PNGase F Digestion of Recombinant S protein produced in HEK or CHO cells

OptSpike1 and OptSpike2 proteins migrated as slightly larger species on SDS-PAGE when expressed in the Expi293F^™^ cells as compared to the ExpiCHO-S^™^ system (**Fig. 2A**), and we hypothesized that this difference in size of the proteins was due to differential N-linked glycosylation associated with Expi293F^™^ and ExpiCHO-S^™^ cells. Heat denatured OptSpike1 was digested with PNGase F, an amidase that cleaves between the innermost GlcNAc and asparagine residues in N-linked glycoproteins, and analyzed by SDS-PAGE. While undigested OptSpike1 produced in Expi293F^™^ cells runs larger than OptSpike1 produced in ExpiCHO-S^™^, the difference in apparent molecular weight is abolished after PNGase F trea™ent (**Fig. 2A**). It is known that the recombinant S proteins are heavily glycosylated, and these results confirm that these proteins exist as distinct N-linked glycoforms when expressed in different mammalian expression systems, as has been previously reported with a number of other glycoproteins [35-37]. Importantly, these differences in glycosylation had no effect on reactivity with sera from convalescent patients (see **Fig. 4B**).

**Figure 2:**
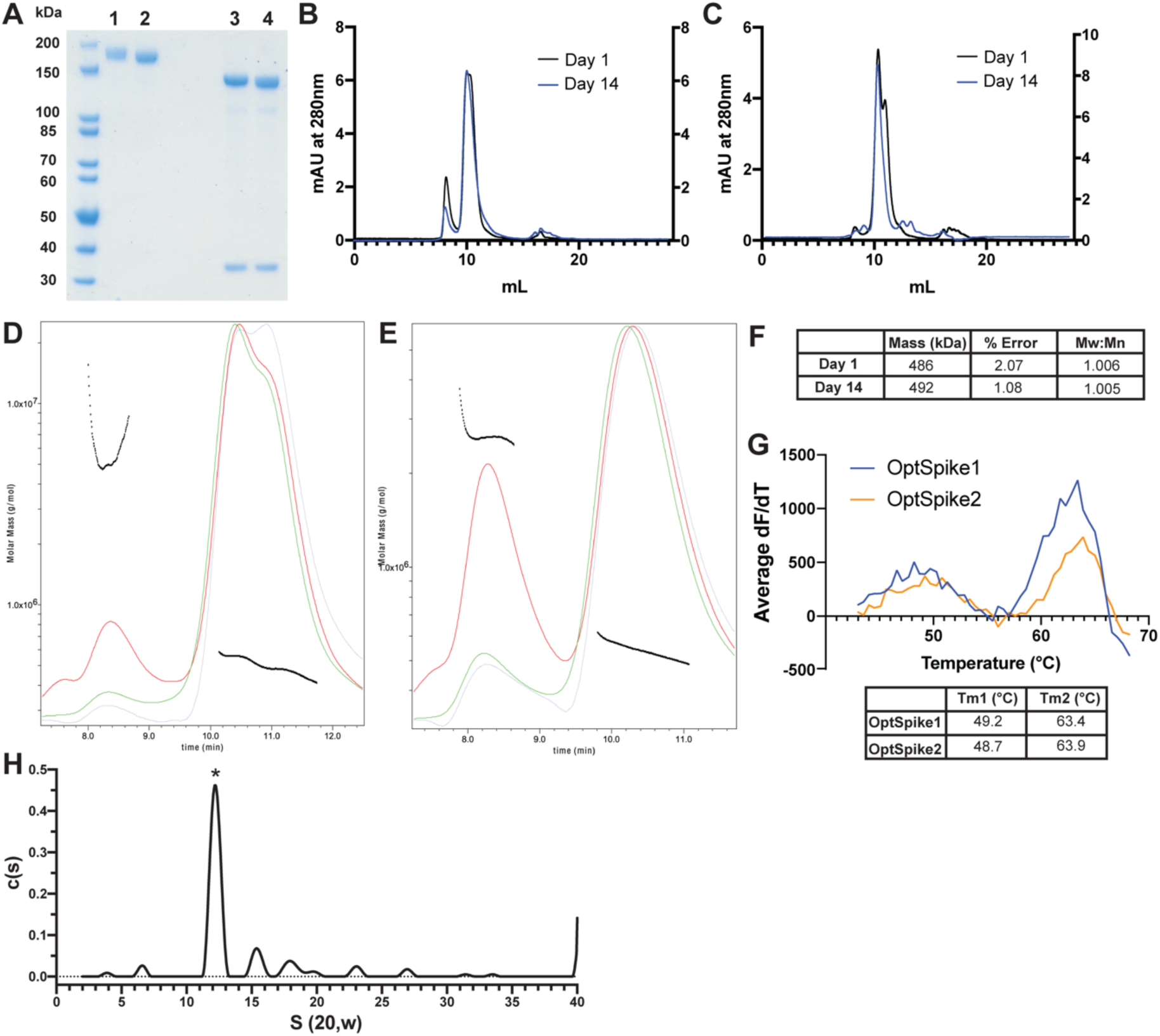
Biophysical Characterization of recombinant SARS-CoV-2 S protein indicates it is stable, uniform, and appropriate molecular mass. **A)** SDS-PAGE analysis of OptSpike1-HEK (lane 1) or OptSpike1-CHO (lane 2) compared to the same protein denatured and treated with PNGase F for 60 minutes (lanes 3 and 4, respectively), lower MW band is PNGase F **B)** Representative SEC traces of OptSpike1-HEK analyzed on a Yarra(tm) 3 µm SEC-4000 LC Column on day 1 post purification (black), and day 14 post purification (blue) **C)** Representative SEC traces of OptSpike1-CHO analyzed on a Yarra(tm) 3 µm SEC-4000 LC Column on day 1 post purification (black), and day 14 post purification (blue) **D)** Representative MALS analysis of OptSpike1-CHO day 1 (red curve: light scattering, green curve: UV280, blue curve: refractive index, black line: Mw) **E)** Representative MALS analysis of OptSpike1-CHO day 14 **F)** Molecular mass of OptSpike1-CHO and PDI (Mw:Mn) determined by MALS **G)** Results of SYPRO Orange thermal denaturation of OptSpike1-CHO and OptSpike2-CHO showing graph of first derivative vs. temperature, and table showing ™1 (left peak), and ™2 (right peak) for each, all experiments are representative of three individual replicates **H)** The c(s) distribution obtained by AUC analysis of OptSpike1-CHO at 4.8 µM total protein concentration. The asterisk denotes the S protein trimeric species. The best fit molecular mass resolved for the trimer is 455 kDa. The best fit ratio of f / f_o_ is 1.63.

**Figure 3:**
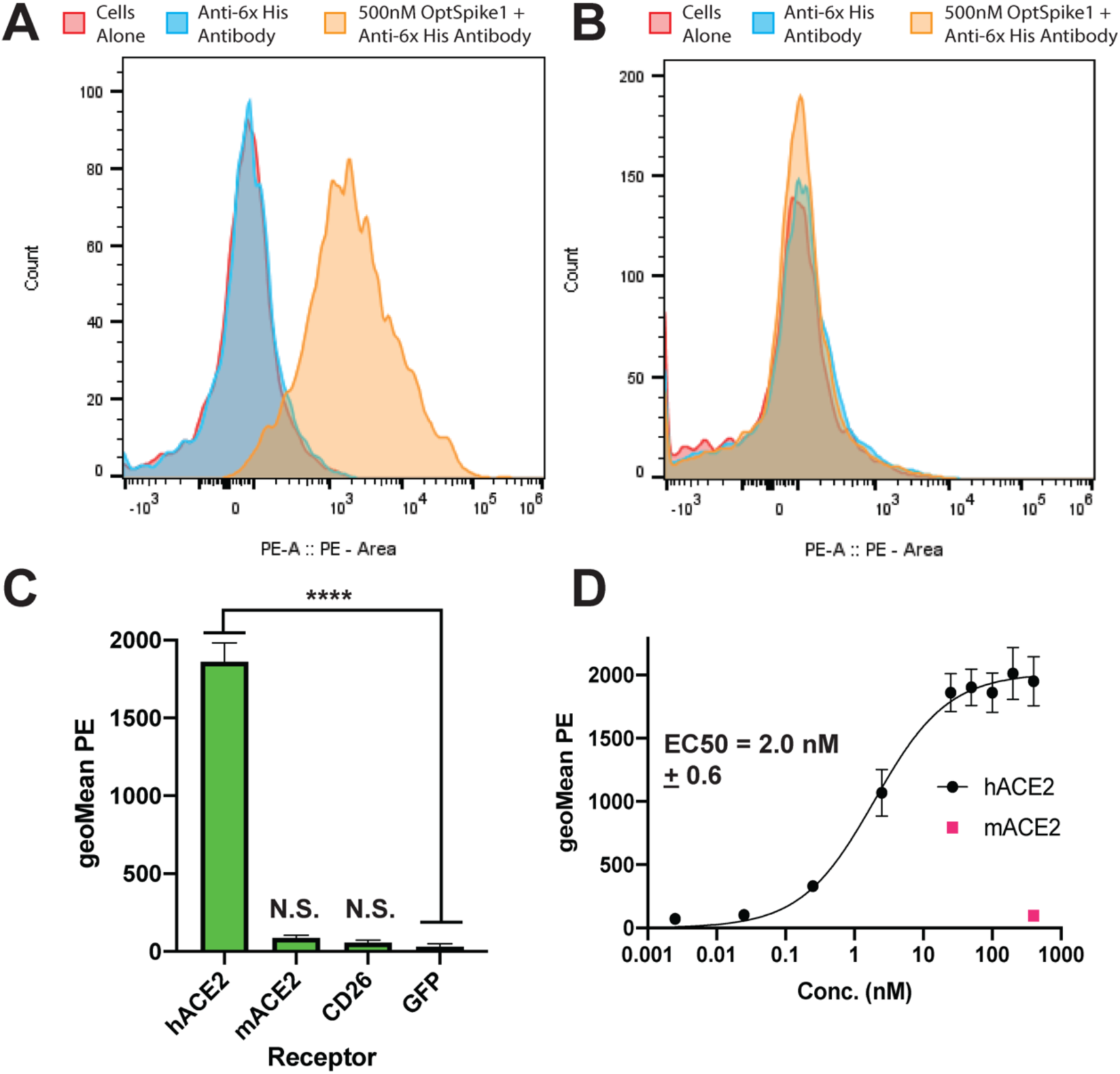
SARS-CoV-2 S Protein Binds to Human ACE2 but not Mouse ACE2 or Human CD26. **A)** Representative flow plot of data quantified in (**C**) showing that OptSpike1-CHO binds to HEK293F cells expressing hACE2, binding was detected using an antibody against the 8x-HIS tag on OptSpike1 **B)** Representative flow plot of data quantified in (**C**) showing that OptSpike1-CHO does not bind to HEK293F cells expressing mACE2, binding was detected using an antibody against the 8x-HIS tag on OptSpike1 **C)** 500 nM OptSpike1-CHO was incubated with cells expressing either human ACE2, mouse ACE2, or human CD26, binding was detected with an anti-HIS antibody and data was acquired by flow cytometry, n=4 **D)** OptSpike1-CHO was titrated on HEK293F cells expressing human ACE2 from .0025-400 nM, binding was quantified with an anti-HIS antibody using flow cytometry, and a binding curve was fit in GraphPad using the equation Y=Bmax*X^h/(Kd^h + X^h), n=4.

**Figure 4:**
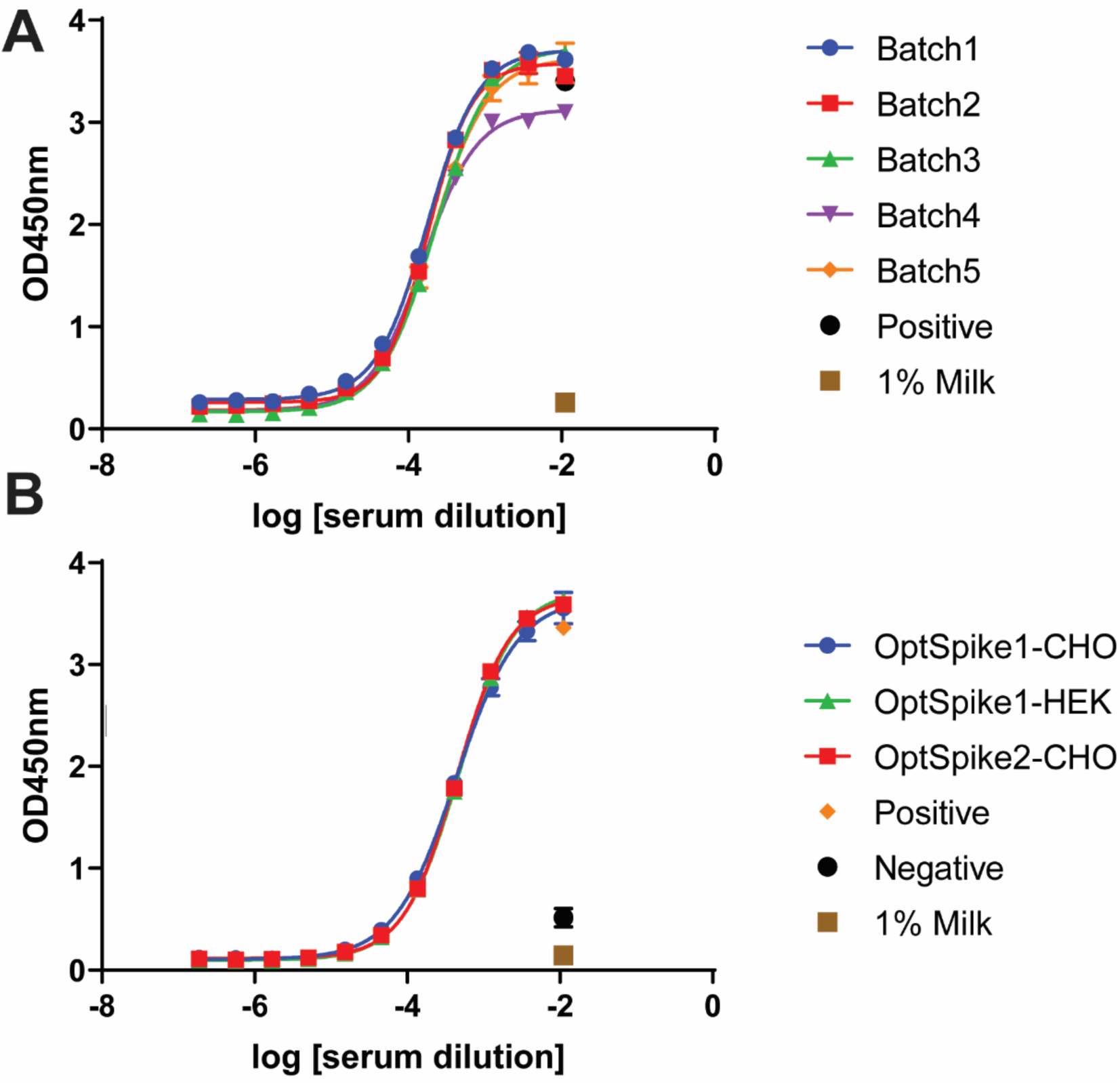
Highly reproducible data in COVID-19 Convalescent serum ELISA experiments using OptSpike1 and OptSpike2: **A)** Five different production batches of OptSpike1-CHO were tested by ELISA for the detection of anti-S IgG antibodies from convalescent COVID-19 patients. Low batch-to-batch variability in OptSpike1 was observed, with no statistically significant differences in EC_50_ values between batches (P>.9999). **B)** Different S protein constructs OptSpike1-CHO, OptSpike1-HEK, and OptSpike2-CHO were tested by ELISA for the detection of anti-S IgG antibodies from confirmed COVID-19 convalescent patients. Very little variability was seen between different constructs and expression cell lines used, with no statistically significant differences in EC_50_ values(P>.9999).

### Size Exclusion Chromatography and Multi Angle Light Scattering (SEC-MALS) analysis of OptSpike1 and OptSpike2

The purified, concentrated OptSpike1 and OptSpike2 proteins were analyzed by FPLC on a Superose^™^ 6 Increase 10/300 GL column (**Fig. S1A-D**), and by HPLC on a Yarra(tm) 3 µm SEC-4000 LC Column (**Fig. 2B-C, Fig. S1E-F**), both of which are appropriate for resolving proteins in the range of, and greater than, the predicted molecular mass (based on amino acid sequence) of ∼420 kDa of trimeric OptSpike1 and OptSpike2. On the Superose^™^ 6 Increase 10/300 GL column OptSpike1 and Optspike2 eluted as a single peak within the included volume, with an apparent molecular mass of ∼670 kDa, and did not form aggregates (**Fig. S1A-D**). Interestingly, analysis on the Yarra(tm) 3 µm SEC-4000 LC Column showed that both OptSpike1 and OptSpike2, expressed in either Expi293F^™^ or ExpiCHO-S^™^ cells, eluted as two partially overlapping peaks (elution times of 10.35 and 11.00 mL) **(Fig. 2B, 2C, Fig. S1E, F**), which could not be resolved on the Superose^™^ 6 Increase 10/300 GL column (**Fig. S1A-D**). Furthermore, after storage at 4°C for fourteen days, the distribution of the elution volume shifted entirely to the faster migrating peak at 10.35 mL (**Fig. 2B-C**). The OptSpike1 produced in ExpiCHO-S^™^ cells was analyzed with Multi Angle Light Scattering (**MALS**) one day after purification, and the polydispersity index (PDI) for species present across both peaks was 1.006, indicating the species across both peaks are uniform with respect to molecular mass (**Fig. 2D, 2F**). The molecular mass of this species was calculated to be 486 kDa ± 2.1% using MALS, which is larger than the predicated molecular mass of 420 kDa (based on the amino acid sequence of trimeric OptSpike1 and not accounting for glycans). The PDI of OptSpike1 on Day 14 (which ran as one peak) was determined to be 1.005, which also indicated that this peak is uniform, and the molecular mass from MALS was 493 kDa± 1.08% (**Fig**.**2E, 2F**). OptSpike-2-HEK, and OptSpike2-CHO exhibited a similar two peak profile when analyzed on the Yarra(tm) 3 µm SEC-4000 LC Column (**Fig. S1E, F**) Together, these data indicate that the existence of closely related species that likely arise from conformational conversion and not from changes in oligomeric state (see below).

### Thermal Denaturation of OptSpike1 and OptSpike2

Differential scanning fluorimetry (**DSF**) was used to assess the relative thermal stabilities of OptSpike1 and OptSpike2 produced in ExpiCHO-S^™^ cells. Analysis of DSF data revealed no discernible difference in melting behavior, with both OptSpike1 and 2 exhibiting two similar melting transitions (OptSpike1 ™_1_ = 49.2° C, ™_2_ = 63.4° C and OptSpike2 ™_1_ = 48.7° C, ™_2_ = 63.9° C) (**Fig. 2G**). This consistent behavior suggests that differences in expression yields of OptSpike1 and Optspike2 are likely not due to inherent differences in protein stability.

### Analytical Ultracentrifugation (AUC) analysis of OptSpike1

To confirm that purified OptSpike1-CHO is a stable trimer and further evaluate the molecular mass, we conducted a series of sedimentation velocity experiments as a function of protein concentration. Six concentrations of S protein were centrifuged spanning a range from 136 nM to 4.8 μM. The distribution of sedimentation coefficients g(s*), and in particular the maxima of the main peak, are invariant over the concentration range examined (**Fig. S1H**). This behavior demonstrates that the protein molecules present in the preparation are stable non-interacting particles. The distribution of species present in a solution of non-interacting particles can be deconvolved using a continuous distribution, c(s), analysis (**Fig. 2H, Fig. S1H**). The dominant peak (68%) in the c(s) distribution has a mass of 455 kDa consistent with the S protein trimer being the dominant protein present in the solution. This calculated molecular mass is comparable to both the predicted molecular mass based on amino acid sequence (420 kDa), and the molecular mass determined by MALS (490 kDa). The best fit ratio of f / f_o_ is 1.63 indicative of significant geometric asymmetry. These data demonstrate that the OptSpike1 is a stable trimer in solution from nM to μM protein concentrations.

### Biochemical Characterization of OptSpike1 interaction with Human ACE2

Purified OptSpike1-CHO protein activity was demonstrated by binding to human ACE2 (**hACE2**), but not mouse ACE2 (**mACE2**) or human CD26 (**hCD26**), the entry receptor for MERS. 500 nM OptSpike1 trimer was incubated with HEK293F cells expressing hACE2, mACE2 or hCD26 —all C-terminally tagged with GFP— and binding was detected by flow cytometry with an anti-His PE labeled antibody recognizing the His_8_ tag on the S protein (see **Fig. S3A**). Expression and cell surface localization of hACE2, mACE2, and hCD26 was confirmed by antibody staining (**Fig. S3E-F, K**). Strong binding to cells expressing hACE2 (**Fig. 3A, 3C**) was observed, but not to cells expressing mACE2 or hCD26 (**Fig. 3B, 3C**), confirming that the S protein does not bind to CD26 (in agreement with previous reports [38]), and confirming specificity of the S protein for hACE2. Titration of hACE2-expression HEK293 cells with recombinant OptSpike1 (0.0025 - 400 nM) yielded an EC_50_ of 2.0 ± 0.6 nM (**Fig. 3D**).

### Screening OptSpike1 for Binding to Members of the Human Secretome

To further evaluate the specificity of the SARS-CoV-2 S protein, OptSpike1-CHO was screened for binding to 900 members (around ∼20%) of the human secretome (from the Ig superfamily, TNFR superfamily, Integrin family, chemokine family, and GPCR family – see **Table S1**). Each member of this library, (tagged with cytosolic GFP to confirm expression – see **Table S1** and **Fig. S3**) was individually transfected into FreeStyle^™^ 293-F cells. Individual transfections were then incubated with 200 nM of OptSpike1 in 96-well plate format and binding was analyzed by flow cytometry with an anti-HIS PE labeled antibody as above. hACE2-GFP expressing cells were included as a positive control on each plate. Integrins were screened both as individual transfections, and as alpha-beta pairs. While strong binding to each replicate of hACE2 was detected, we did not detect binding to any other members of the library (**Fig. S3, Table S1**), including proteins suggested to be targets of S protein binding, including CD147 [39, 40], Siglec 9 and 10 [41], as well as the host cell receptors for other coronaviruses, including CEACAM1, which is the receptor for Murine Coronavirus[42], and CD26 and CEACAM5, which are both receptors for MERS [43]. Expression and cell surface presentation of these putative receptors was validated using monoclonal antibodies (**Fig. S3I-L**).

### OptSpike1 and OptSpike2 can be used reproducibly and interchangeably in COVID-19 convalescent serum ELISA

Enzyme-linked immunosorbent assays (**ELISA**s) are commonly used in clinical settings to detect the presence of viral antibodies. Therefore, we assessed the reactivity of serum antibodies from one COVID-19 convalescent patient toward multiple independent preparations the S ectodomain proteins (**Fig. 4A**). We analyzed the EC_50_ and standard error by one-way ANOVA and found that there is no statistical significance in EC_50_ values when different production batches of OptSpike1 were used as the target (**Fig. 4A**, p>.9999), or when the different S protein constructs OptSpike1-CHO, OptSpike2-CHO, and OptSpike1-HEK were used as the target **(Fig. 4B**, p>.9999). These data demonstrate that the expression and purification protocols reported herein consistently yield OptSpike1 with reproducible behavior in ELISAs detecting anti-S IgG antibodies. Additionally, we have shown that reactivity of serum antibodies from convalescent patients toward OptSpike1 and OptSpike2 is not distinguishable.

### Development of a COVID-19 Multi-antigen Protein Array

Protein microarray technology allows for the high-throughput, multiplexed screening of numerous parameters within a single experiment [44]. To simultaneously and rapidly screen the serum from convalescent COVID-19 patients against multiple SARS-CoV-2 antigens at once, we developed a COVID-19 multi-antigen protein array presenting purified S protein, the RBD of the S protein (**Fig. S2C**) and Nucleocapsid protein (**Fig. S2A**,**B**,**D**) of SARS CoV-2 (printed with a Marathon Argus piezoelectric printer from Arrayjet). This multi-antigen array was challenged with either convalescent sera from COVID-19 patients that had been previously diagnosed using RT-PCR and ELISA tests, or serum from control individuals that had been collected prior to the SARS-CoV-2 outbreak in November of 2019. Seropositivity of COVID-19 patients was confirmed by ELISA (**Fig. S5**). In total, serum from eight individuals was screened. Individuals 2, 3, 5, 6, 7 are confirmed SARS-CoV-2 positive. Individuals 1, 4 and 8 had serum collected before SARS-CoV-2 was reported (**Fig. 5** and **Fig. S4**).

**Figure 5:**
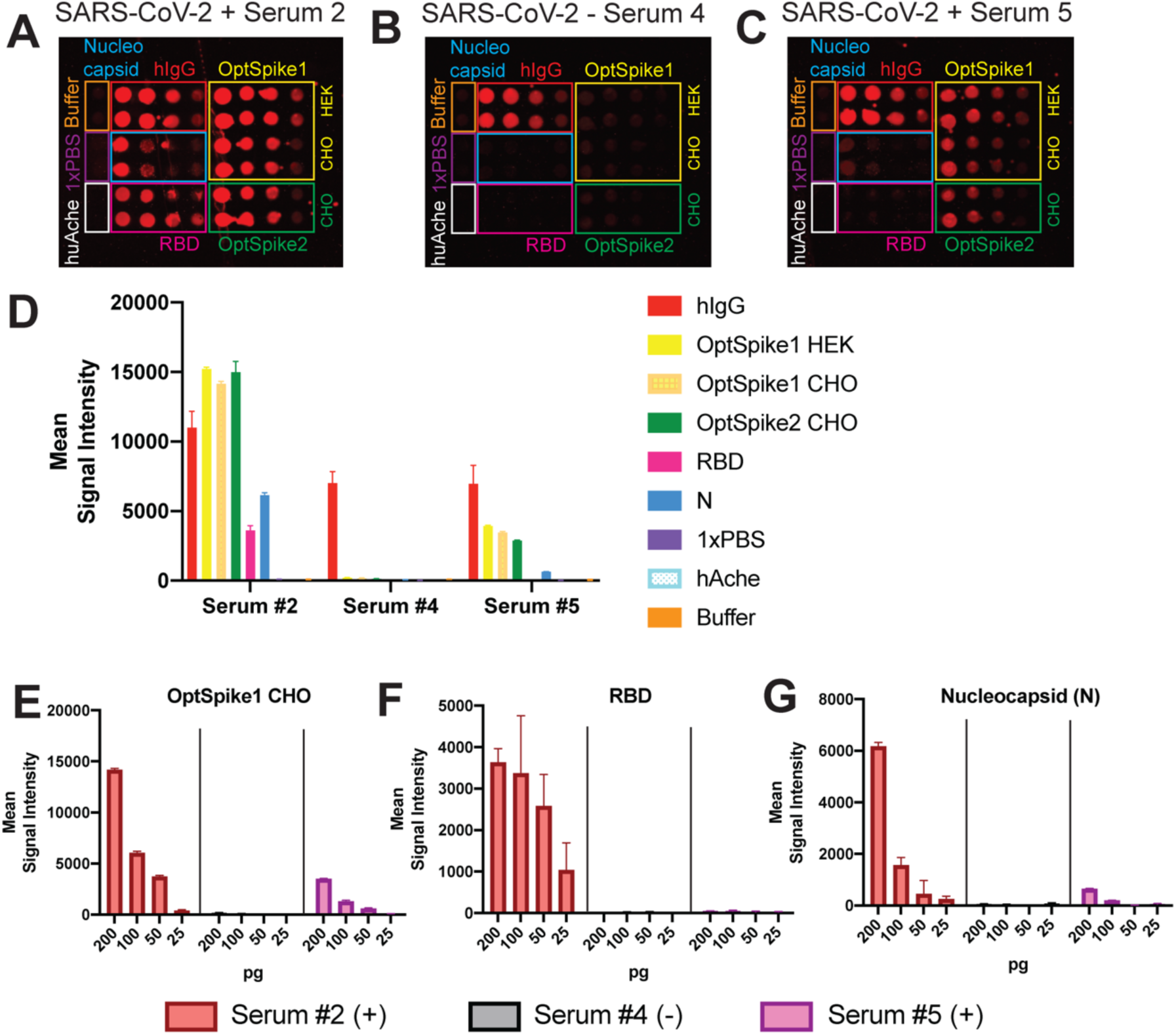
SARS-CoV-2 Multi-antigen Protein Array. **A)** Representative image of antigen detection from SARS-CoV-2 positive serum 2. Serum 2 detects all of the SARS-CoV-2 antigens printed on the protein array: S protein (OptSpike1 and 2), Receptor Binding Domain of S protein (RBD) and Nucleocapsid protein (N). (**B**) Representative image of screening with SARS-CoV-2 negative serum 4. Serum 4 is negative for all SARS-CoV-2 antigens printed and only positive control, human IgG1 (hIgG1) is detected. (**C**) Representative image of antigen detection from SARS-CoV-2 positive serum 5. Serum 5 does not detect all of the SARS-CoV-2 antigens printed on the protein array and only detects S protein and N protein. All protein arrays contain negative buffer and 1xPBS controls as well as negative protein controls (huAche). **E)** Quantifications of serum 2, 4 and 5 results with titrations of OptSpike1 produced in ExpiCHO-S cells from 200pg to 25pg. **F)** Quantifications of serum 2, 4 and 5 results with titrations of RBD from 200pg to 25pg. **G)** Quantifications of serum 2, 4 and 5 results with titrations of N protein from 200pg to 25pg.

Each array was printed with sixteen identical subarrays. A range of target protein concentrations were spotted (25, 50, 100 and 200 pg per spot) and each serum was screened in duplicate (**Fig. 5A-C, Fig. S4A-B**). Recombinant human IgG1, which will be recognized by the anti-human-647 labeled secondary antibody, was included in each array as a positive control for protein printing. Negative controls (human acetyl cholinesterase (**huAche**), printing buffer and PBS) were used to determine nonspecific binding and background correction for data analysis. All confirmed COVID-19 patients had strong antibody responses to the S protein in our multi-antigen protein array, which was in good agreement with the S protein ELISAs (**Fig. S4D-F, Fig. S5**). Additionally, all confirmed COVID-19 patient sera tested positive for N protein in our protein multi-antigen array (**Fig. S4H**). Antibodies against the RBD were detected in serums 2, 3, 6, and 7. However, antibodies against the RBD from patients 5 (which additionally had relatively lower detectable levels of antibodies against the S and N protein) could not be detected in this assay (**Fig. 5D-G, Fig. S4 C-H**). These data demonstrate the ability of the COVID-19 Multi-antigen Protein Array to simultaneously analyze antibody responses to multiple antigens in a high-throughput format and again validate the use of our recombinant S protein in an antigen detection platform. Understanding variable antibody responses to the spectrum of SARS-CoV-2 antigens will substantially impact our understanding of population-wide immune responses to COVID-19.

### Cryo-EM Structure of the SARS-CoV-2 S Protein produced in ExpiCHO-S^™^ cells

Cryogenic electron microscopy (**cryo-EM**) single particle reconstruction was conducted to assess the tertiary and quaternary arrangement of the S protein expressed in the HEK and CHO systems (**Table S2**). Aliquots of purified OptSpike1 protein from each expression host (1mg/mL in 50mM Tris and 250mM NaCl, pH 8.0) were applied to cryo-EM grids, blotted and plunge-frozen in liquid ethane at LN_2_ temperature. Micrographs collected on a Titan Krios at 300kV, using a K3 camera, were aligned and motion corrected followed by CTF estimation and 2D classification as described in the methods. Particle picking, 3D refinement, and reconstruction yielded high quality maps for samples arising from both HEK and CHO production (**Fig. S6, Table S2**). Using the OptSpike1-CHO material, we obtained a 3.22 Å resolution reconstruction of the symmetrical trimer in the closed conformation (all three RBD in the down position) (**Fig. 6A-B, Fig. S6, Table S2**). Similarly, the OptSpike1-HEK material yielded a 3.44 Å reconstruction of the symmetrical trimer in the closed conformation (**Fig. S6, Table S2**). Rigid body refinement and manual adjus™ent in COOT followed by RSR refinement against the reconstructed maps with PHENIX [45] yielded models with good stereochemistry and validation metrics (**Table S2, Fig. 6, Fig. S6A-F**). Comparison of the molecular envelopes from the current work with the recently deposited cryo-EM structure of the closed state of the SARS Cov-2 SPIKE protein (**PDB: 6VXX**) showed excellent agreement. For example, structural alignment of the trimeric CHO-produced S protein coordinates with those from 6VXX resulted in a core R.M.S.D. of 0.63Å for 2889 aligned Cα pairs. Likewise, comparison of the coordinates for the HEK and CHO coordinates from the current work aligned with R.M.S.D. of 0.60Å over 2696 aligned Cα pairs. This is consistent with proper tertiary folding and trimeric quaternary organization of both samples [19].

**Figure 6:**
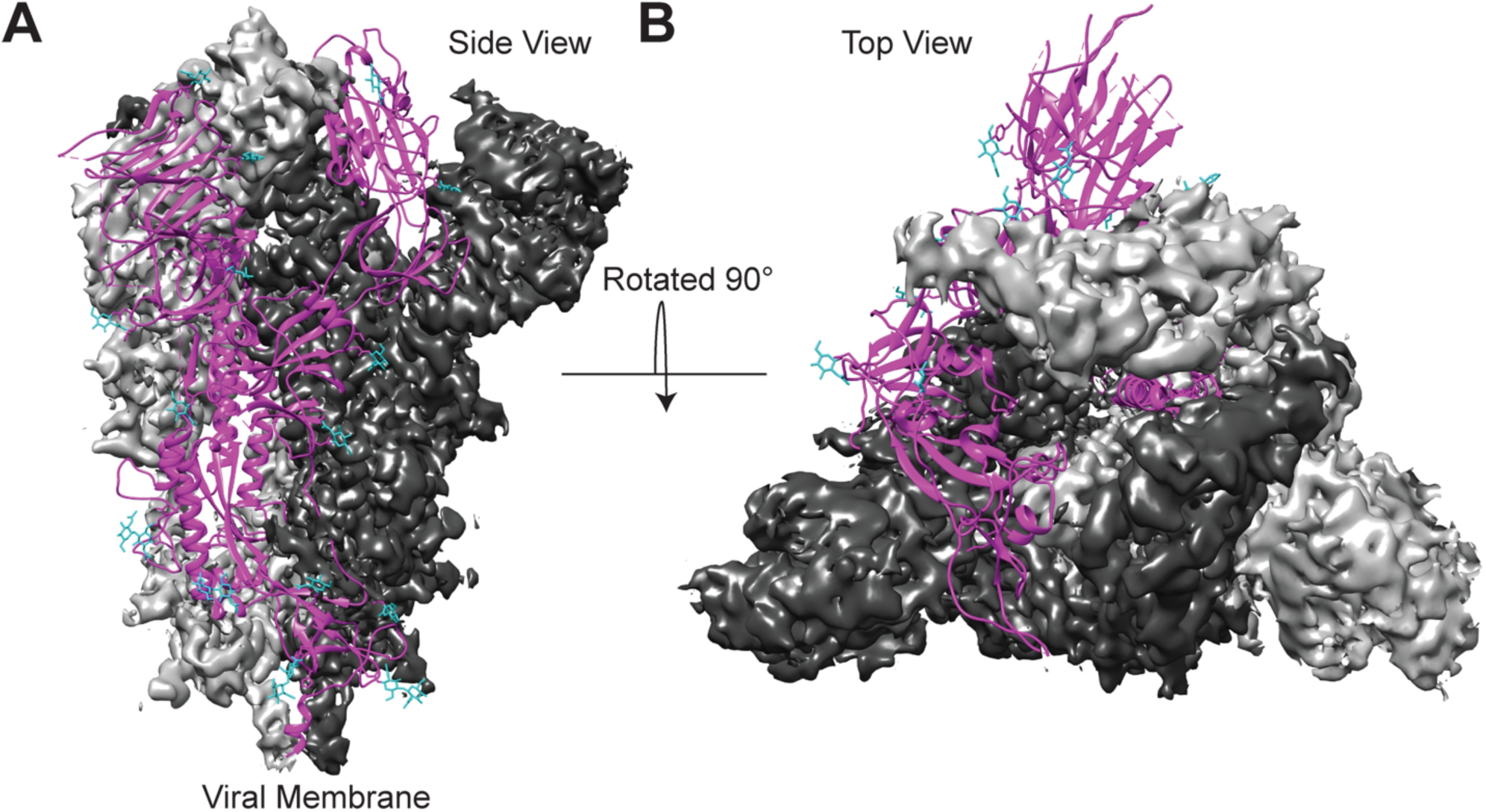
Cryo-EM structure of OptSpike1-CHO in the closed state. **A)** Side view of the SARS-CoV-2 OptSpike1 trimer in the prefusion conformation. **B)** Top view of the SARS-CoV-2 OptSpike1 trimer in the prefusion conformation. Two protomers are displayed with the cryo-EM density maps (dark grey and light grey) and the third protomer is displayed as a ribbon structure (magenta) with glycans represented on the ribbon structure (cyan).

## DISCUSSION

To effectively address the COVID-19 global health crisis, reproducibility in biological and clinical studies will be critical. Efficient and uniform production of key biological reagents is essential for the generation of reliable data in the coming months and years. Herein, we describe our protocols (including SOP) for the optimized production of two recently reported recombinant forms of the SARS-CoV-2 S protein. We show that using the ExpiCHO-S^™^ expression system, recombinant S protein can be reproducibly produced in both high quantities and high quality. Our reported yield of 28 mg/L for OptSpike1, and 18 mg/L for OptSpike2, exceed those previously reported (OptSpike1: .5 mg/L in HEK293F cells [7], OptSpike2: 5 mg/L in Expi293F^™^ cells, and .5 mg/L in insect cells [13]). To validate protein quality, we employed analytical size exclusion chromatography, multi angle light scattering, thermal denaturation and analytical ultracentrifugation, and demonstrated that these recombinant S proteins have similar biophysical properties and antigenicity.

Interestingly, we found that all preparations consistently exhibited different behavior when examined on two different analytical size exclusion columns. These proteins eluted as a single peak on the Superose^™^ 6 Increase 10/300 GL column (agarose-based resin), but eluted as two overlapping peaks on the Yarra(tm) 3 µm SEC-4000 (silica-based resin). We also noted a time-dependent evolution, as the distribution of peaks on the Yarra(tm) 3 µm SEC-4000 moved entirely to the faster migrating species. Furthermore, MALS analysis demonstrated that there was no mass difference between the two overlapping peaks and AUC analysis of OptSpike1-CHO confirmed that the predominant species in solution was a stable trimer. We propose a modest time-dependent structural alteration of the S protein, involving two states with distinct physico-chemical properties, resulting in differential interactions with the silica-based Yarra(tm) 3 µm SEC-4000 resin. This proposal is consistent with reported cryo-EM structures displaying the trimeric S protein exists in both a closed conformation with all RBDs pointing downwards (**PDB: 6VXX**) and an open conformation with one RBD pointing upwards (**PDB: 6VSB** and **6VYB**) [7, 19]. Future work will be required to fully define the nature and functional consequences of the multiple solution species observed for the S protein.

We demonstrate that the protein purified here is well suited for structural studies by determining the cryo-EM structures of OptSpike1-HEK at 3.44Å resolution and OptSpike1-CHO at 3.22Å. These structures are nearly identical to each other and are in good agreement with the closed form observed in previously reported structures of the S protein (which were produced in HEK293F cells) [7, 19]. Interestingly, the structures determined from protein expressed in mammalian cells differ from a recently reported cryo-EM structure of wild type S protein produced from High Five^™^ insect cells. This structure appeared more compact overall and the model could not accommodate the presence of the SD1 domain [46]. Whether these disparities are the consequence of differences in expression hosts (e.g., glycosylation or other processing functions) or the lack of stabilizing mutations, remains an open question, although RBD recognition and antigenicity was preserved.

Both OptSpike1 and OptSpike2 proteins exhibit comparable thermal denaturation profiles with two similar melting transitions (**Fig. 2G**). This finding is consistent with the thermal denaturation profile of wild type S protein purified from High Five^™^ insect cells, despite the overall structural differences [46]. Additional work will be required to assign these transitions to specific domains or modules within the full length COVID-19 spike protein.

Additionally, we examined the receptor-binding specificity of the SARS-CoV-2 S protein. We demonstrate strong binding between the S protein and hACE2, but not mACE2, as has been previously reported [8, 47]. Furthermore, we screened recombinant OptSpike1 against a sub-library of 900 members of the human secretome that were expressed on HEK-293 cells and did not identify any additional interactions (**Fig. S3C**). This library included many proteins that others have suggested may bind to the SARS-CoV-2 S protein, including CD147, Siglec 9, Siglec 10, and proteins that other coronaviruses use for viral entry, including CD26 (MERS), CEACAM5 (MERS), and CEACAM1 (Murine Coronavirus). [41, 42, 48, 49]. CD147 is of particular interest because of the ongoing clinical trial that aims to treat SARS-CoV-2 with Meplazumab (a humanized anti-CD147 antibody) [39]. Importantly, for this subset of proteins, we conducted antibody staining to explicitly evaluate cell surface localization (**Fig. S3I-L**). While all members of this subset could be detected by antibody staining, and are therefore properly localize to the cell surface, we were not able to detect binding to the S protein. It is important to note that the assay we employed represents only one possible format for detecting protein-protein interactions (recombinant S protein binding to cell surface displayed receptor: **Fig. S3A**). Because we did not directly recapitulate the experiments that originally detected interactions between the S protein and CD147 (SPR, ELISA, Co-IP[40]) or Siglec 9 and 10 (ELISA [41]), it is possible that the S protein binds to these receptors in a way that is undetectable in this assay, or that there will be cell-specific differences in presentation of the putative receptors (i.e., the requirement an unknown co-receptor not expressed on HEK293 cells, etc.). It is important to note that anti-CD147 Abs were reported to block COVID-19 infection in Vero E6 cells, which also express ACE2 [50], highlighting the potential importance of cell-specific differences in receptor presentation and recognition. Further investigations are warranted to evaluate these potential interactions. Finally, as our screening efforts only included ∼20% of the human secretome, it is important to continue to search for and evaluate alternate interactions, which might reveal additional mechanisms of viral entry for SARS-CoV-2.

We additionally demonstrate that these recombinant S proteins can be used interchangeably and reproducibly in two different assays screening for a human serum response to S protein (ELISA: **see Fig. 4**, and protein microarray: see **Fig. 5, S4** and below). Validating that both S proteins can be utilized in a serum ELISA screen is important because both OptSpike1 and OptSpike2 are being widely utilized for serology testing at different clinical sites. These data collectively confirm that the strategy for antigen preparation described here can be used reliably, and that the antigenicity of OptSpike1 and OptSpike2 are comparable, regardless of cell line used (HEK vs. CHO: **Fig. 5B**). These data are critical for interpreting clinical result from various institutions, which are using a wide array of different serological tests to detect anti-S antibodies (for a complete list of antibody tests approved for use by the FDA, please see [51]). Consequently, additional versions of the S protein that are being used clinically should be analyzed similarly, so as to further ensure clinical reproducibility and compatibility of different antibody serology tests.

We developed a protein microarray platform to simultaneously analyze serum for antibodies against the S protein, as well as recombinant RBD and N protein. Microarray results from a small cohort of serum from 5 seroreactive SARS-CoV-2 patients and serum from 3 individuals collected before November of 2019 revealed that detectable anti-S antibody titer correlated well with results of serological ELISAs (**Fig. S4** and **S5**), and additionally demonstrated the simultaneous detection of antibodies against the S protein, the N protein, and the RBD of the S protein. It is important to note the limitations of the current platform. In particular, we stress that at present this is a qualitative approach, as a number of variables can impact the ability to detect antibody reactivity, including patient titers for specific antigens and the relative affinities of an antigen-specific pool. Furthermore, it should be appreciated that the current platform is programmed to detect the capture of IgG antibodies; thus, the resultant signal (or lack thereof) for a particular antigen could be the consequence of prevalences between different isotypes (IgA, IgD, IgE, IgG, IgM), which are known to evolve during the course of infection and subsequent resolution. Although currently focused on three antigens, this platform can be readily expanded to study differential antibody responses to different SARS-CoV-2 antigens and subdomains of those antigens amongst individuals, which are actively being investigated by others [52-55]. We are currently working to include not only other antigens from SARS-CoV-2 (E protein, M protein, etc.), but also antigens from other coronaviruses that may be cross reactive with SARS-CoV-2 antibodies. The analysis of antibody reactivity to multiple SARS-CoV-2 and related antigens will provide broad insight into the humoral immune response to SARS-CoV-2.

Collectively, we provide standards and metrics for high quality protein reagents that can yield comparable clinical, biological and structural data as we continue to combat the global health crisis caused by SARS-CoV-2.

## ACKNOWLEDGEMENTS

This work was supported by the NIH (R01 CA198095 to S.C.A., R01-AI125462 to J.R.L., 5R01-GM129350 to M.B., U19AI142777 and R01AI132633 to K.C., R01AI143453 and R01AI123654 to L.P., and R21AI141367 to J.P.D). Some of this work was performed at the Simons Electron Microscopy Center and National Resource for Automated Molecular Microscopy located at the New York Structural Biology Center, supported by grants from the Simons Foundation (SF349247), NYSTAR, and the NIH National Institute of General Medical Sciences (GM103310) with additional support from the National Center for CryoEM Access and Training (NCCAT) supported by the NIH Common Fund Transformative High Resolution Cryo-Electron Microscopy program (U24 GM129539,) and NY State Assembly Majority. A.J.N. was supported by a grant from the NIH National Institute of General Medical Sciences (NIGMS) (F32GM128303). M.E.D. is a Latin American Fellow in the Biomedical Sciences, supported by the Pew Charitable Trusts. J.H.L, R.H.B.III., and R.J.M. were partially supported by NIH training grant 2T32GM007288-45 (Medical Scientist Training Program) at Albert Einstein College of Medicine. L.P. was additionally supported by a grant from the Mathers Foundation.

We thank Jason McLellan for providing the stabilized spike plasmid (OptSpike1), and Arvin Dar, Sarah Pegno, and Zaigham Khan for sharing OptSpike2 plasmid. We also thank Daija Bobe at SEMC and Al Tucker at Einstein for technical support, Phaneendra Duddempudi for useful discussion. We also thank Bridget Carragher and Clint Potter for assistance and input in cryo-EM work.

## DATA AVAILABILITY

The accession numbers for the EM maps, models, and raw movie stack for the SPIKE structure from protein produced in CHO cells reported in this paper are EMD-22078, PDB Entry 6×6P, and EMPIAR-10433.

