## Supplemental Figures and Appendix 1 for "Characterization of the SARS-CoV-2 S Protein: Biophysical, Biochemical, Structural, and Antigenic Analysis"

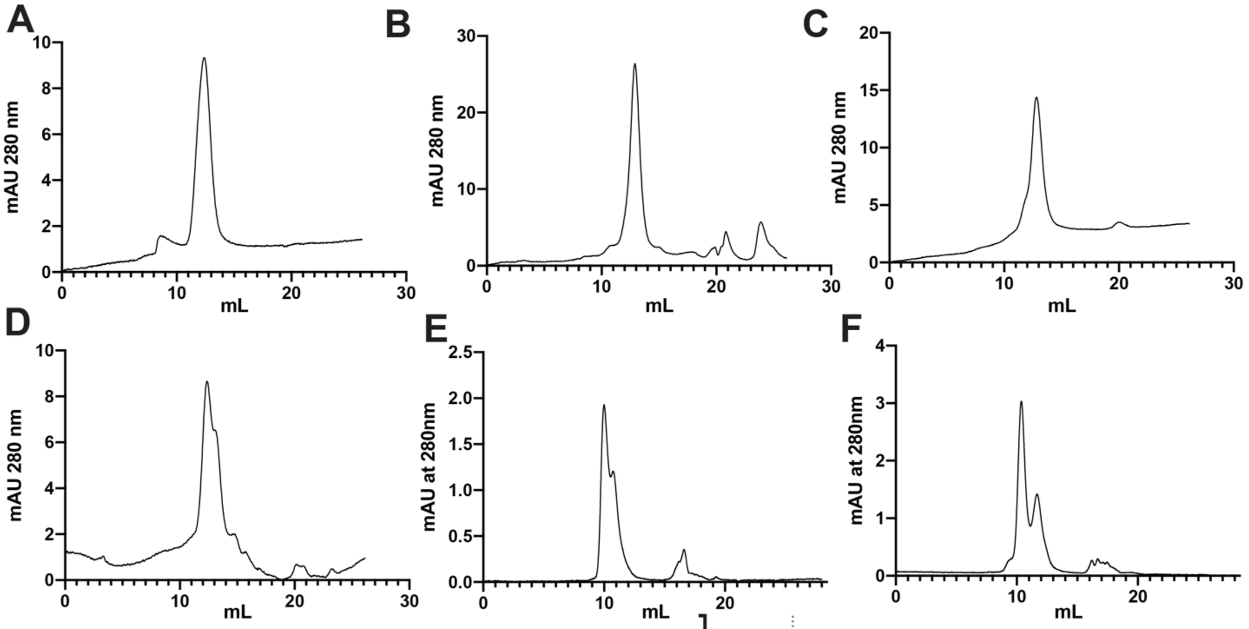


G

| Cell System | Expression Length | Incubation Temp | Yield |
|---|---|---|---|
| ExpiCHO-S Standard | 4 days | 37°C | 6.4 mg/mL |
| ExpiCHO-S High Titer | 4 days | 32°C | 4.9 mg/mL |
| ExpiCHO-S Standard | 8 days | 37°C | 17.1 mg/mL |
| ExpiCHO-S High Titer | 8 days | 32°C | 20.9 mg/mL |
| FreeStyle 293-F | 4 days | 37°C | 0.53 mg/mL |
| FreeStyle 293-F | 6 days | 37°C | 1.33 mg/mL |

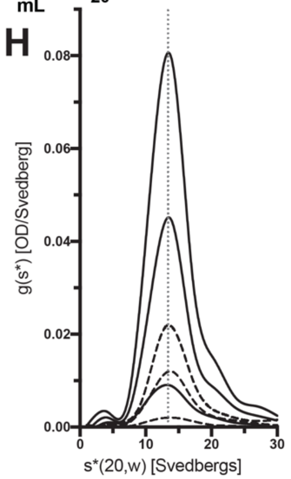


**Supplemental Figure 1: Quality Control of S Proteins.** Analytical size exclusion chromatography of purified S proteins on a Superose™ 6 Increase 10/300 GL column in the following order: **A)** Expi293F produced OptSpike1. **B)** ExpiCHO-S produced OptSpike1. **C)** Expi293F produced OptSpike2. **D)** ExpiCHO-S produced OptSpike2. S proteins were also analyzed via HPLC on a Yarra™ 3 $\mu$m SEC-4000 LC Column and while OptSpike1 analysis is in figure 2 we are including the OptSpike2 analysis in the following order: **E)** Expi293F produced OptSpike2. **F)** ExpiCHO-S produced OptSpike2. **G)** Table indicating expression differences between ExpiCHO standard titer and ExpiCHO max titer protocol and in FreeStyle 293-F for OptSpike1 **H)** Smoothed time – derivative g(s*) distribution plots of the six AUC sedimentation runs that were conducted on OptSpike1-CHO. The solid lines depict the data acquired scanning at 280 nm. The dashed lines depict the data acquired scanning at 230 nm with the O.D. values scaled to the 280 nm data by their relative extinction coefficients. The vertical dotted grey line highlights the invariance of the S protein trimer peak maxima with protein concentration; this invariance as a function of protein concentration is a hallmark of a non-interacting system. The g(s*) distributions were generated using the program DCDT+ by John Philo v2.5.1.

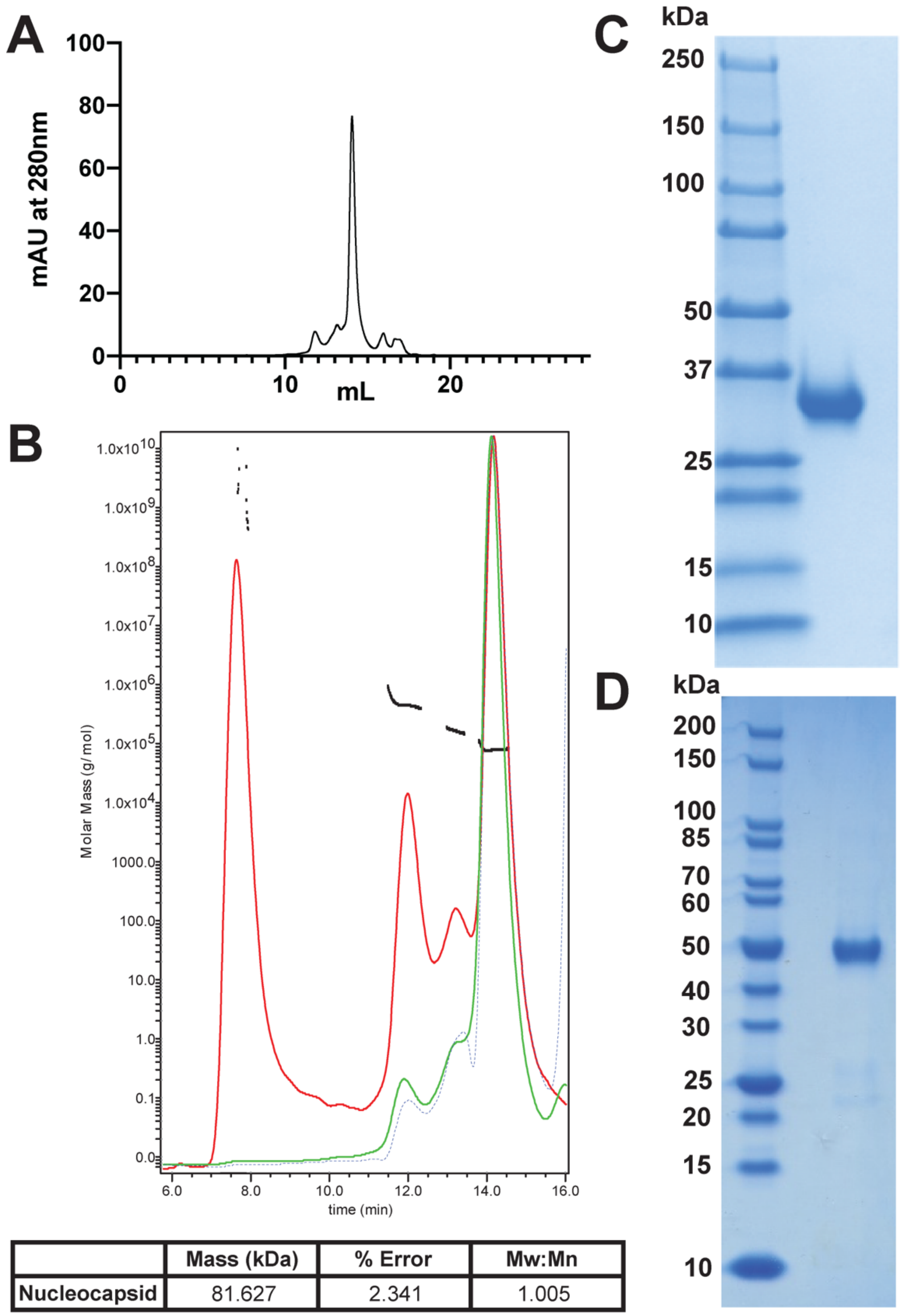


| | Mass (kDa) | % Error | Mw:Mn |
|---|---|---|---|
| Nucleocapsid | 81.627 | 2.341 | 1.005 |

**Supplemental Figure 2: Production and Characterization of the SARS-CoV-2 N and RBD Proteins. A)** Analytical size exclusion chromatography of purified N Antigen run on a Superose™ 6 Increase 10/300 GL column. **B)** Representative SEC MALS data for N antigen (red curve: light scattering, green curve: UV280, blue curve: refractive index, black line: Mw:Mn). **C)** Purified RBD produced in FreeStyle 293-F cells. **D)** SDS-PAGE analysis of N antigen purification.

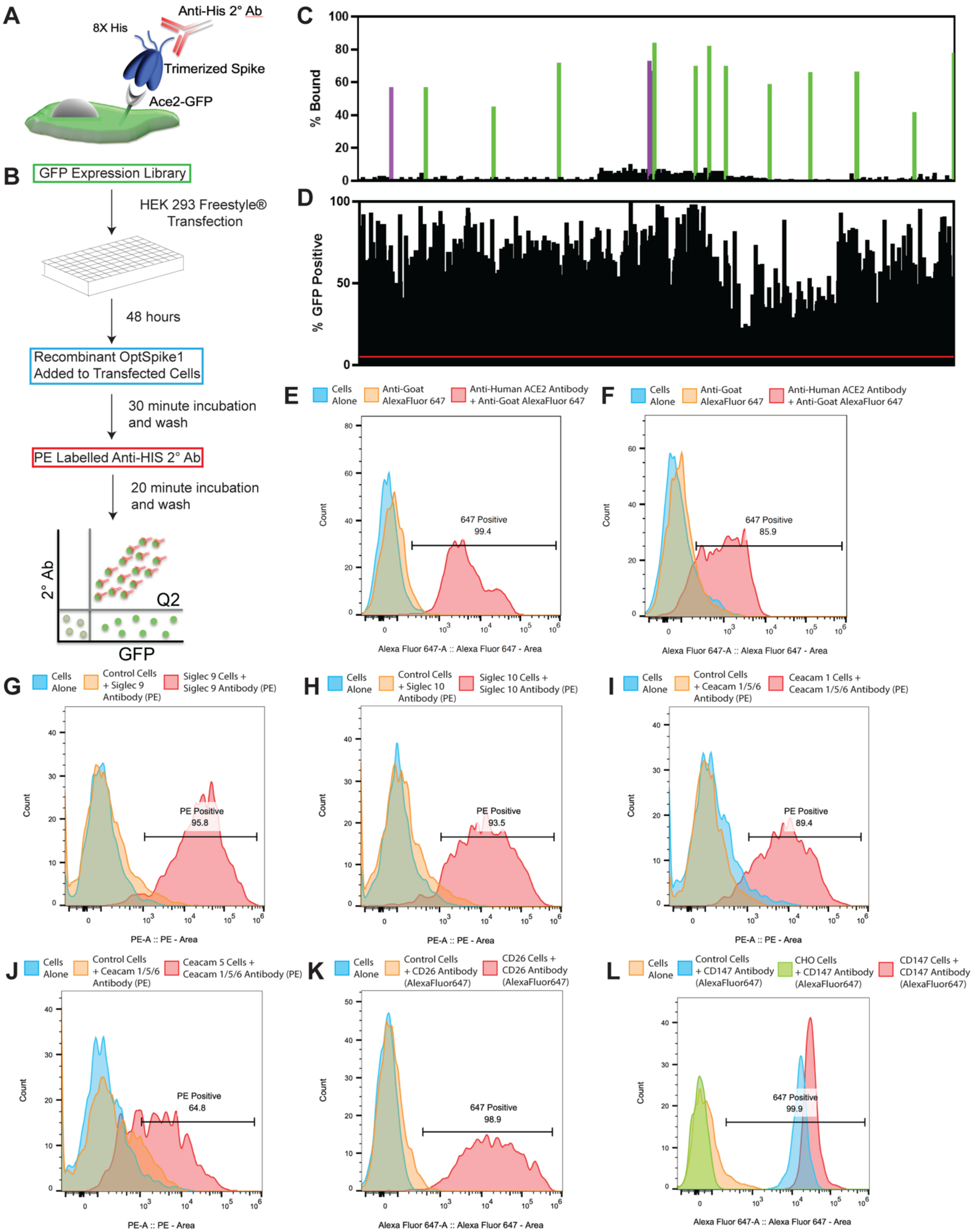
A
8X His
Anti-His 2° Ab
Trimerized Spike
Ace2-GFP
B
GFP Expression Library
HEK 293 Freestyle® Transfection
48 hours
Recombinant OptSpike1 Added to Transfected Cells
30 minute incubation and wash
PE Labelled Anti-HIS 2° Ab
20 minute incubation and wash
2° Ab
Q2
GFP
C
% Bound
D
% GFP Positive
E
Cells Alone
Anti-Goat AlexaFluor 647
Anti-Human ACE2 Antibody + Anti-Goat AlexaFluor 647
647 Positive 99.4
Count
Alexa Fluor 647-A :: Alexa Fluor 647 - Area
F
Cells Alone
Anti-Goat AlexaFluor 647
Anti-Human ACE2 Antibody + Anti-Goat AlexaFluor 647
647 Positive 85.9
Count
Alexa Fluor 647-A :: Alexa Fluor 647 - Area
G
Cells Alone
Control Cells + Siglec 9 Antibody (PE)
Siglec 9 Cells + Siglec 9 Antibody (PE)
PE Positive 95.8
Count
PE-A :: PE - Area
H
Cells Alone
Control Cells + Siglec 10 Antibody (PE)
Siglec 10 Cells + Siglec 10 Antibody (PE)
PE Positive 93.5
Count
PE-A :: PE - Area
I
Cells Alone
Control Cells + Ceacam 1/5/6 Antibody (PE)
Ceacam 1 Cells + Ceacam 1/5/6 Antibody (PE)
PE Positive 89.4
Count
PE-A :: PE - Area
J
Cells Alone
Control Cells + Ceacam 1/5/6 Antibody (PE)
Ceacam 5 Cells + Ceacam 1/5/6 Antibody (PE)
PE Positive 64.8
Count
PE-A :: PE - Area
K
Cells Alone
Control Cells + CD26 Antibody (AlexaFluor647)
CD26 Cells + CD26 Antibody (AlexaFluor647)
647 Positive 98.9
Count
Alexa Fluor 647-A :: Alexa Fluor 647 - Area
L
Cells Alone
Control Cells + CD147 Antibody (AlexaFluor647)
CHO Cells + CD147 Antibody (AlexaFluor647)
CD147 Cells + CD147 Antibody (AlexaFluor647)
647 Positive 99.9
Count
Alexa Fluor 647-A :: Alexa Fluor 647 - Area

**Supplemental Figure 3: Screening OptSpike1 Against 20% of the Human Secretome. A)** Schematic illustrating binding of purified OptSpike1 to ACE2-expressing cells and detection of binding by anti-His antibody. **B)** Depiction of the workflow for screening our GFP expression library of the human secretome. **C)** Results from screening 840 transmembrane proteins from the IG, TNFR, integrin, GPCR and chemokine families of the human secretome. HEK293F cells expressing members of the human secretome were incubated with 200 nM OptSpike1 in 96-well plate format, ACE2 was included in every plate as a positive control (green bars). S protein binding to cells was detected using an anti-HIS antibody (PE), thus FC receptor expressing cells were detected (purple bars). Binding was assessed by flow cytometry and data was analyzed by sub gating GFP positive events, and then PE positive events. **D)** GFP expression profile of cells analyzed in **(C)**, constructs that did not have more than 5% GFP positive events (red line) were excluded from analysis. **E)** Validation of surface localization of human ACE2 expressing cells using a polyclonal anti-human ACE2 antibody. **F)** Validation of surface localization of mouse ACE2 expressing cells using the same antibody as in **(E)**. **G)** Validation of surface localization of human Siglec 9 using PE-labeled anti-Siglec 9 antibody **H)** Validation of surface localization of human Siglec 10 using PE-labeled anti-Siglec 10 antibody **I)** Validation of surface localization of human Ceacam1 using PE-labeled anti-Ceacam1/5/6 (anti-CDD66a/c/e) antibody **J)** Validation of surface localization of human Ceacam5 using PE-labeled anti-Ceacam1/5/6 (anti-CDD66a/c/e) antibody **K )** Validation of surface localization of human CD26 using Alexa647-labeled anti-CD26 antibody **L)** Representative flow plots showing that HEK293F cells endogenously express high levels of CD147, and that expression is increased upon transfection with CD147-GFP. CHO cells do not express CD147 and were used as a negative control. Staining was performed using an Alexa647-labeled anti-CD147 antibody.

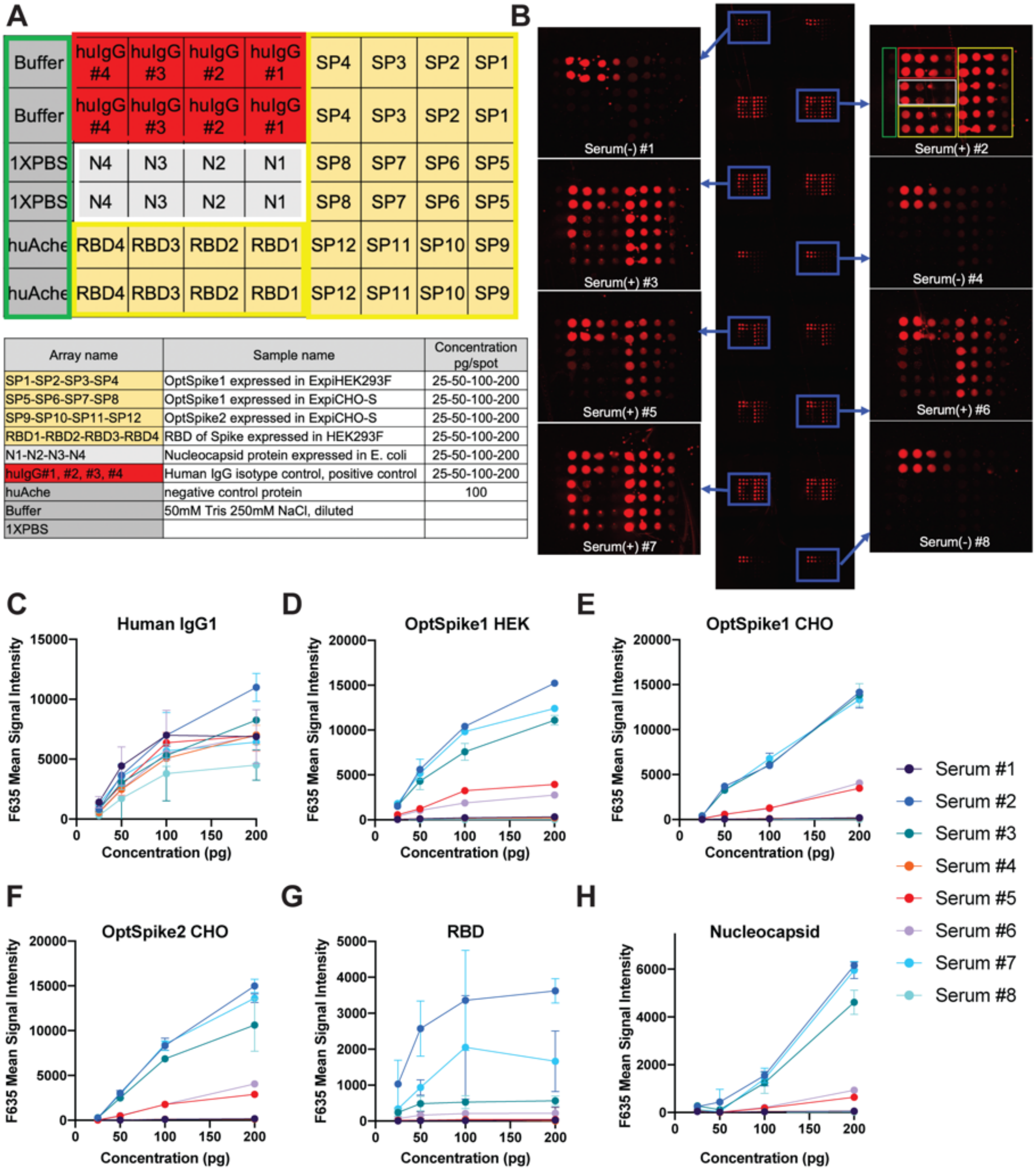


| Array name | Sample name | Concentration pg/spot |
|---|---|---|
| SP1-SP2-SP3-SP4 | OptSpike1 expressed in ExpiHEK293F | 25-50-100-200 |
| SP5-SP6-SP7-SP8 | OptSpike1 expressed in ExpiCHO-S | 25-50-100-200 |
| SP9-SP10-SP11-SP12 | OptSpike2 expressed in ExpiCHO-S | 25-50-100-200 |
| RBD1-RBD2-RBD3-RBD4 | RBD of Spike expressed in HEK293F | 25-50-100-200 |
| N1-N2-N3-N4 | Nucleocapsid protein expressed in E. coli | 25-50-100-200 |
| hulgG#1, #2, #3, #4 | Human IgG isotype control, positive control | 25-50-100-200 |
| huAche | negative control protein | 100 |
| Buffer | 50mM Tris 250mM NaCl, diluted | |
| 1XPBS | | |

**Supplemental Figure 4**: **SARS-CoV-2 Multi-antigen Protein Array**. **A)** Diagram depicting layout of proteins printed in (**B**) and table indicating proteins printed at various concentrations. SARS-CoV-2 antigens printed: Spike Protein (SP), Receptor Binding Domain of Spike protein (RBD) and Nucleocapsid protein (N). Buffer controls were printed and in addition, a positive control protein human IgG isotype control (huIgG) and a negative control protein (huAche) were printed. **B**) Representative images of protein array post detection: after incubation with serum samples, antibodies against immobilized proteins are detected by Alexa Fluor 647-labeled secondary anti-human IgG antibody. Each array is printed with 16 identical sub-arrays to screen each serum sample in duplicate. Convalescent serum from COVID-19 patients is indicated as (+) whereas control human serum is indicated as (-). The sub-arrays 1, 4 and 8 were used to screen COVID19 (-) serum samples; sub-arrays 2, 3, 5, 6 and 7 were used to screen COVID-19 (+) serum samples. **C)** Human IgG1 positive control titrations for serum 1-8. **D)** OptSpike1 HEK titrations for serum 1-8. **E)** OptSpike1 CHO titrations for serum 1-8. **F)** OptSpike2 CHO titrations for serum 1-8. **G)** RBD titrations for serum 1-8. **H)** Nucleocapsid titrations for serum 1-8.

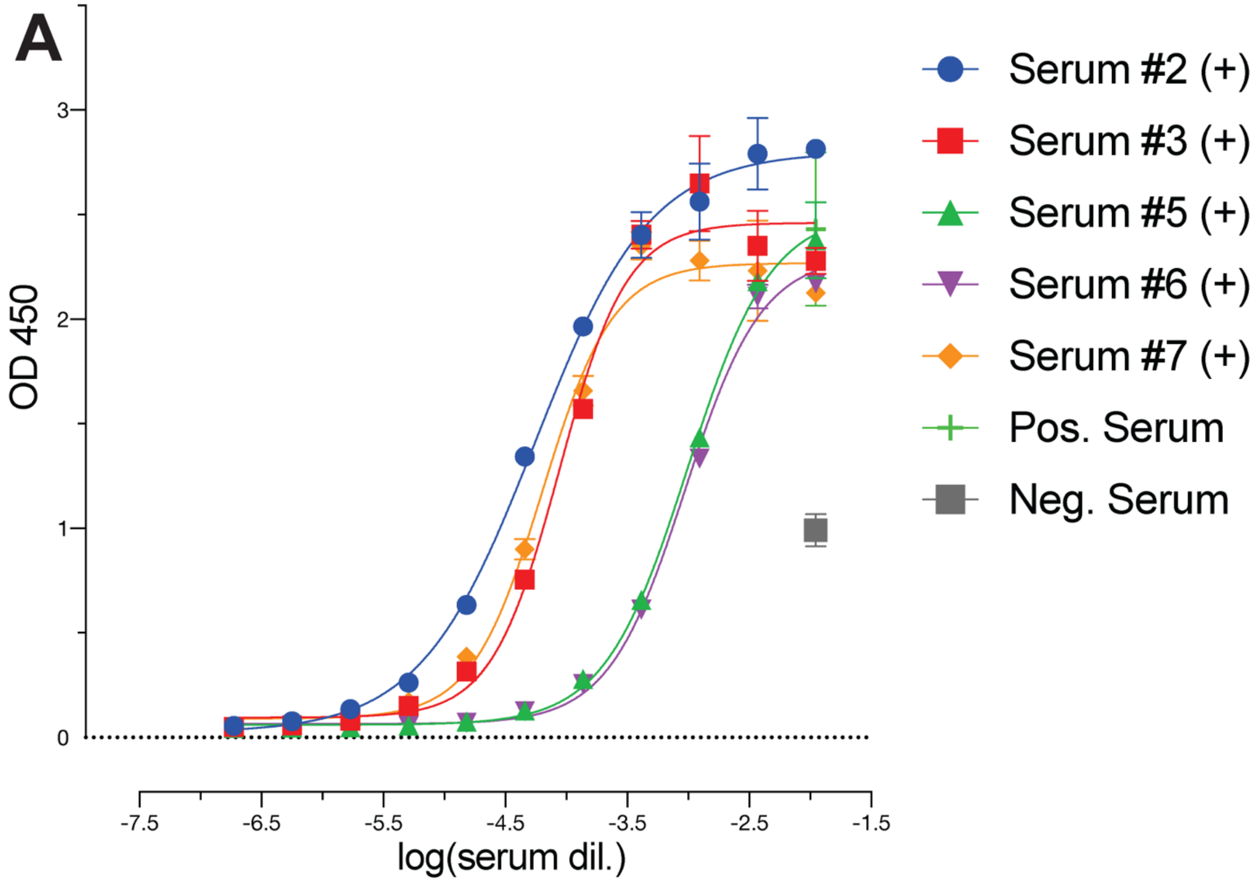


**Supplemental Figure 5: Representative ELISAS Confirming S Protein reactivity of Positive Serum.** Serum from confirmed COVID-19 positive patients used in COVID-19 multi-antigen array was tested by ELISA for the detection of anti-S IgG antibodies.

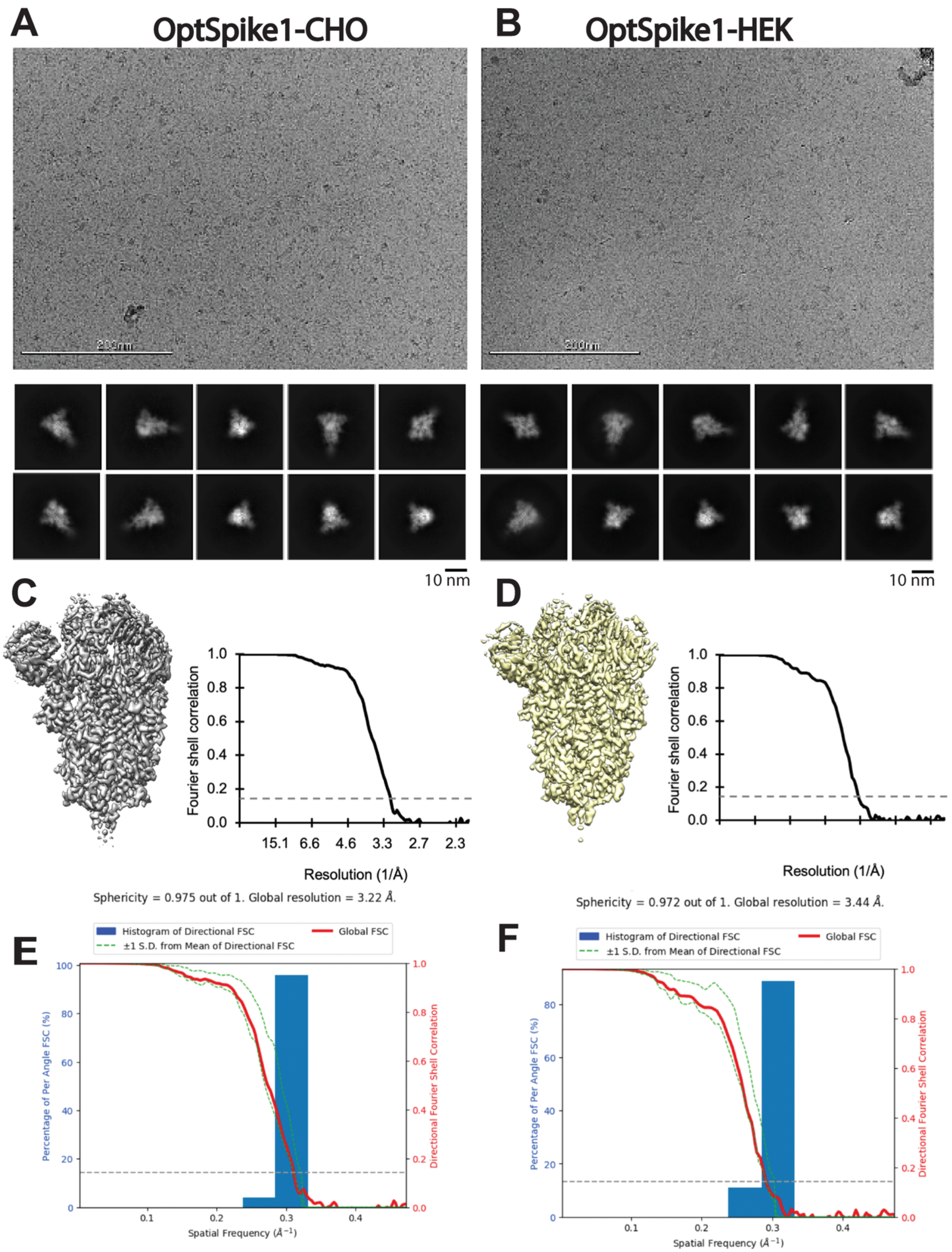

A
OptSpike1-CHO
B
OptSpike1-HEK
10 nm
10 nm
C
D
Fourier shell correlation
1.0
0.8
0.6
0.4
0.2
0.0
15.1
6.6
4.6
3.3
2.7
2.3
Resolution (1/Å)
Sphericity = 0.975 out of 1. Global resolution = 3.22 Å.
Sphericity = 0.972 out of 1. Global resolution = 3.44 Å.
E
F
Histogram of Directional FSC
±1 S.D. from Mean of Directional FSC
Global FSC
Percentage of Per Angle FSC (%)
Directional Fourier Shell Correlation
Spatial Frequency (Å⁻¹)

**Supplemental Figure 6: Representative micrographs, class averages, isosurface representations, and Directional FSC profile of OptSpike1-CHO and OptSpike1-HEK.** Representative micrograph and 2D class averages of OptSpik1-CHO (**A**) and OptSpike1-HEK (**B**), respectively. The 3D reconstruction and Fourier shell correlation (FSC) curve (with the 0.143 cut off annotated by the grey dotted line of (**C**) OptSpike1-CHO in grey at 3.22Å and (**D**) OptSpike1-HEK in yellow at 3.44Å, respectively. 3D FSC plot showing the directional FSC of (**E**) OptSpik1-CHO and (**F**) OptSpike1-HEK, respectively using the Remote 3DFSC Processing Server (https://3dfsc.salk.edu/, http://dx.doi.org/10.1038/nmeth.4347).

**Table S2.** Cryo-EM data collection, refinement, and validation statistics

| | SARS-Cov2 SPIKE CHO | SARS-Cov2 SPIKE HEK |
|---|---|---|
| **Data collection and processing** | | |
| Microscope | TFS Titan Krios | TFS Titan Krios |
| Detector | Gatan K3/GIF (20eV) | Gatan K3/GIF (20eV) |
| Magnification | 81,000× | 81,000× |
| Voltage (kV) | 300 | 300 |
| Electron exposure ($e^-/Å^2$) | 66.5 | 66.5 |
| Defocus range (μm) | 0.8–2.5 | 0.8–2.5 |
| Pixel size (Å) | 1.058 | 1.058 |
| Symmetry imposed | C3 | C3 |
| Initial particle images (no.) | 75,582 | 99,154 |
| Final particle images (no.) | 54,395 | 54,066 |
| Map resolution (Å) | 3.22 | 3.44 |
| FSC threshold | 0.143 | 0.143 |
| Initial model used (PDB code) | 6VXX | 6VXX |
| Model resolution (Å) | 3.22 | 3.44 |
| FSC threshold | 0.143 | 0.143 |
| Model resolution range (Å) | ∞−3.22 | ∞–3.44 |
| Map sharpening *B*-factor ($Å^2$) | −86 | −95 |
| **Model composition** | | |
| Non-hydrogen atoms | 24,615 | 23,694 |
| Protein residues | 3,051 | 2,916 |
| Ligand molecules | 57 | 63 |
| *B*-factors ($Å^2$) | 120 | 100 |
| R.M.S. deviations | | |
| Bond lengths (Å) | 0.012 | 0.010 |
| Bond angles (°) | 1.041 | 1.046 |
| Validation | | |
| MolProbity score (percentile) | 2.69 | 2.71 |
| Clashscore (percentile) | 24.29 | 24.48 |
| Poor rotamers (%) | 3.04 | 4.81 |
| Ramachandran plot | | |
| Favored (%) | 92.93 | 95.46 |
| Allowed (%) | 7.07 | 4.54 |

| | | |
|---|---|---|
| Disallowed (%) | 0.00 | 0.00 |

**Appendix 1.** SARS-CoV2 S "Spike" protein Expression and Purification SOP

**SARS-CoV2 Expression in ExpiCHO-S Cells**

**Required Equipment**
Class II biological safety cabinet
CO2 Incubator with shaker (we recommend either: Infors HT Multitron **OR** Climo Shaker ISF4-X)
Hemocytometer
Pipet-Aid
Micropipettes
Refrigerator at 4°C (+/- 1°C)

**Required Materials and Reagents**
pCAGGs SARS-COV2 stabilized S protein plasmid** (See below)
ExpiCHO-S Cells (Thermo Fisher Scientific, A29127)
ExpiCHO expression medium (Thermo Fisher Scientific, A29100)
ExpiFectamine CHO Transfection Kit (Thermo Fisher Scientific, A29129)
OptiPRO Serum-Free Medium (Gibco, Thermo Fisher Scientific, 12309-050)
1 L non-baffled, vented flask, sterile (Fisher, PBV1000)
Trypan blue solution, 0.4 % (Gibco #15250-06)
1.5 mL Eppendorf tubes (Denville #C2170)
Polypropylene sterile conical tubes
- 15 mL (Denville Scientific #C1018P or equivalent)
- 50 mL (Denville Scientific #C1060P or equivalent)

Sterile, serological pipettes
- 10mL (Falcon #357551 or equivalent)
- 25 mL (Falcon #357535 or equivalent)

Micropipette tips
- 20 $\mu$L barrier tips (Denville Scientific #P1121 or equivalent)
- 200 $\mu$L barrier tips (Denville Scientific #P1122 or equivalent)
- 1000 $\mu$L barrier tips (Denville Scientific #P1126 or equivalent)

**Procedure**

<u>Seeding ExpiCHO cells (Day -1)</u>

1. Cells are passaged every 3-4 days at a density of 4-6 million cells/mL and incubated in an orbital shaking incubator at 37°C and 125 RPM with 8% CO2.
2. Count cells by taking a 20 uL aliquot of suspended cells and adding an equivalent volume of Trypan blue. Load onto hemocytometer to count cells and ensure cells are >95% viable.

3. Seed cells 24 hours before transfection at 3 million cells/mL in a 200 mL volume in a 1 L vented flask.

Transfecting ExpiCHO cells (Day 0) – Example of a 200mL ExpiCHO-S Transfection

1. Transfections are done as per manufacturer's instruction. Determine viable cell density by trypan blue staining.
2. Dilute the cells to final density of 6 million viable cells/mL with fresh ExpiCHO Expression Medium, pre-warmed to 37°C. Swirl the flasks gently to mix the cells.
3. In a 50 mL conical tube, add 200 ug of plasmid DNA to 8 mL of OptiPro. Invert to mix.
4. In a 15 mL conical tube, add 7.4 mL of Optipro and 640 mL of Expifectamine CHO reagent. Invert to mix.
5. Add OptiPro/Expifectamine mixture to OptiPro/Plasmid DNA mixture. Invert tube to mix.
6. Incubate at room temperature for 5 minutes.
7. Add 16 mL of OptiPro/DNA/Expifectamine mixture to 200 mL cell suspension in 1L flask.
8. Incubate at **37°C** in shaking incubator (125 RPM with 8% CO2)

**The ExpiCHO-S Expression System offers three different expression protocols, we utilize two of these protocols to express SARS-CoV-2 S protein:**

Adding Feed and Enhancer to Transfected ExpiCHO Cells (Day 1)- Example of feeding cells following the **Max Titer** Protocol:

1. Feed cells as per manufacturer's instructions.
2. In a 50 mL conical tube, add **32 mL** of ExpiCHO Feed. Add **1.2 mL** of ExpiCHO Enhancer and invert tube.
3. Add Feed/Enhancer mixture to cells.
4. Incubate at **32°C** in shaking incubator (125 RPM with 8% CO2) until harvest (**Day 12**).

Adding Feed to Transfected ExpiCHO Cells (Day 5)- Example of feeding cells following the **Max Titer** Protocol:

1. In a 50 mL conical tube, add **32 mL** of ExpiCHO Feed
2. Add Feed mixture to cells.
3. Incubate at **32°C** in shaking incubator (125 RPM with 8% CO2) until harvest (**Day 12**).

**SARS-CoV-2 Spike Glycoprotein Purifications under Native Conditions**

**Required Equipment**
Sorvall ST 40R centrifuge or equivalent

**Required Materials and Reagents**
500mL Conical tube (Corning Cat#431123) or conical tube of appropriate volume
SIGMAFAST Protease Inhibitor Cocktail tablets, EDTA-Free (Sigma Cat#S8830-20TAB Lot#SLCC0143)
10mL Serological Pipette (Sarstedt Cat#86.1254.001)
Ni-NTA purification resin (Goldbio Cat# H-350-100 Lot# 1363.031520A) **OR**
His60 $Ni^{2+}$ superflow resin (Takara Cat: 635664)
Econo-Pac Chromatography Columns (Bio-Rad Cat#732-1011)
1X Native Binding Buffer (50mM Tris HCl pH8.0, 250mM NaCl)
1X Native Wash Buffer (50mM Tris HCl pH8.0, 250mM NaCl, 10mM Imidazole)
1X Native Elution Buffer (50mM Tris HCl pH8.0, 250mM NaCl, 500mM Imidazole)
1X Storage Buffer (50mM Tris HCl pH8.0, 150mM NaCl)
100K concentrator (ThermoFisher Cat # 88533)
1 mL Syringe (NORM-JECT Tuberkulin Cat#4010-200V0 Lot#19H05C8)
Millex- GV 0.22$\mu$m Filter Unit (Cat# SLGV004SL Lot#R4CA46158)
Nalgene™ Rapid-Flow™ Sterile Disposable Filter Units with PES Membrane (ThermoFisher Cat#524-0020)
2 M ArgCl, pH 6.5
Slide-A-Lyzer™ G2 Dialysis Cassettes (Thermo Fisher 87725)

**NOTE**: Filter all buffer solutions and distilled water through 0.22$\mu$m filter units prior to use (unless being used for dialysis)

Cell Supernatant Harvest (Day 12)

1. Transfer cell suspension into 500mL Conical tubes and centrifuge at 500g for 10 min to pellet cells
2. Transfer supernatant to clean tubes and further centrifuge at 2000g for 10min to clarify
3. Pass clarified supernatant through Nalgene Rapid Flow Disposable Filter Unit with PES Membrane
4. Supplement with 1 tablet protease inhibitors per 100mL supernatant
5. Adjust supernatant to 50 mM ArgCl using a 2 M stock solution

Ni-NTA or His60 $Ni^{2+}$ Purification bead preparation

1. Thoroughly re-suspend Nickel resin to create a slurry
2. For every **1 L** of clarified supernatant use **50 mL** of suspended Nickel resin, and distribute into chromatography columns
    a. Snap off column spout to allow drainage of flow-through

3. Allow Nickel resin to drain
4. Wash Nickel resin by adding 2 resin volume distilled water to each column and allow to drain
5. Wash Nickel resin by adding 2 resin volume 1X Native Binding Buffer to each column and allow to drain
6. Cap column and resuspend resin in 1 resin volume of Native Binding Buffer

Nickel Batch Binding

1. Resuspend resin into slurry
2. For every 1 Liter of clarified supernatant, add 100 mL of resuspended resin (50 mL of resin)
3. Place on a shaker or rotating apparatus for 3 hours at 4˚C

SARS-CoV-2 Spike Purification

1. Following batch binding, distribute supernatant containing Nickel resin equally amongst chromatography columns
2. Allow all supernatant to flow through the columns
3. Wash resin in each column 2 times with 10 resin volumes 1X Native Wash Buffer
4. Add 2 resin volumes of 1X Native Elution Buffer to each column and collect flow-through containing SARS-CoV-2 Spike glycoprotein in 100K concentrator

**If performing an ELISA with the SARS-CoV-2 Spike protein, then Buffer Exchange is sufficient for final purification step:**

Buffer Exchange, Protein Concentration and Quantitation and Storage

1. Centrifuge Eluant collected in 100k concentrator at 1100g, at 4˚C in 8-minute intervals
2. In between 8-minute intervals, ensure protein dispersity by either inverting concentrator, or pipetting up and down
3. Check protein concentration in between centrifugation steps using nanodrop (do not allow protein to become more concentrated than 1 mg / ml, and continue until protein is at desired volume for dialysis
4. Dialyze protein overnight against 1X Storage Buffer at 4 ˚C in appropriate dialysis cassette (use 500 mL storage buffer for 35 mL dialysis cassette)
5. Store protein at 4˚C for short-term periods, or flash freeze in liquid N2 and store at -80˚C for long-term periods
6. For longer 4˚C storage periods, adjust storage buffer to include 100 mM ArgCl, 10% glycerol (if appropriate for downstream applications)

**If higher quality spike protein is necessary (i.e., for structural studies), then size exclusion chromatography should be performed after nickel affinity and before dialysis:**

Size Exclusion Chromatography

**Required Equipment:**

Size exclusion chromatography column and FPLC: We recommend HiLoad 16/600 Superdex 200 (GE)

1. Equilibrate column in 1X storage buffer, for longer 4˚C storage periods, adjust storage buffer to include 100 mM ArgCl, 10% glycerol (if appropriate for downstream applications)
2. After concentrating to appropriate volume, purify protein on HiLoad 16/600 Superdex 200, collecting all fractions
3. Spike protein will elute with an apparent MW of 670 kDa.
4. Collect appropriate fractions, avoiding any aggregate species present, and concentrate and store as described above. After SEC, do not concentrate to more than .8 mg / mL unless absolutely required for downstream applications

**SARS-CoV2 Stabilized Spike AA Sequence**

MFVFLVLLPLVSSQCVNLTTRTQLPPAYTNSFTRGVYYPDKVFRSSVLHSTQDLFLPFFS
NVTWFHAIHVSGTNGTKRFDNPVLPFNDGVYFASTEKSNIIRGWIFGTTLDSKTQSLLIV
NNATNVVIKVCEFQFCNDPFLGVYYHKNNKSWMESEFRVYSSANNCTFEYVSQPFLMD
LEGKQGNFKNLREFVFKNIDGYFKIYSKHTPINLVRDLPQGFSALEPLVDLPIGINITRFQT
LLALHRSYLTPGDSSSGWTAGAAAYYVGYLQPRTFLLKYNENGTITDAVDCALDPLSET
KCTLKSFTVEKGIYQTSNFRVQPTESIVRFPNITNLCPFGEVFNATRFASVYAWNRKRISN
CVADYSVLYNSASFSTFKCYGVSPTKLNDLCFTNVYADSFVIRGDEVRQIAPGQTGKIA
DYNYKLPDDFTGCVIAWNSNNLDSKVGGNYNYLYRLFRKSNLKPFERDISTEIYQAGST
PCNGVEGFNCYFPLQSYGFQPTNGVGYQPYRVVVLSFELLHAPATVCGPKKSTNLVKN
KCVNFNFNGLTGTGVLTESNKKFLPFQQFGRDIADTTDAVRDPQTLEILDITPCSFGGVS
VITPGTNTSNQVAVLYQDVNCTEVPVAIHADQLTPTWRVYSTGSNVFQTRAGCLIGAEH
VNNSYECDIPIGAGICASYQTQTNSPGSASSVASQSIIAYTMSLGAENSVAYSNNSIAIPTN
FTISVTTEILPVSMTKTSVDCTMYICGDSTECSNLLLQYGSFCTQLNRALTGIAVEQDKNT
QEVFAQVKQIYKTPPIKDFGGFNFSQILPDPSKPSKRSFIEDLLFNKVTLADAGFIKQYGD
CLGDIAARDLICAQKFNGLTVLPPLLTDEMIAQYTSALLAGTITSGWTFGAGAALQIPFA
MQMAYRFNGIGVTQNVLYENQKLIANQFNSAIGKIQDSLSSTASALGKLQDVVNQNAQ
ALNTLVKQLSSNFGAISSVLNDILSRLDPPEAEVQIDRLITGRLQSLQTYVTQQLIRAAEIR
ASANLAATKMSECVLGQSKRVDFCGKGYHLMSFPQSAPHGVVFLHVTYVPAQEKNFTT
APAICHDGKAHFPREGVFVSNGTHWFVTQRNFYEPQIITTDNTFVSGNCDVVIGIVNNTV
YDPLQPELDSFKEELDKYFKNHTSPDVLGDISGINASVVNIQKEIDRLNEVAKNLNESLI
DLQELGKYEQGSGYIPEAPRDGQAYVRKDGEWVLLSTFLGRSLEVLFQGPGHHHHHHH
HSAWSHPQFEKGGGSGGGGSGGSAWSHPQFEK

Foldon Trimerization Motif

PreScission Site

His Tag

StrepII Tag

<u>NT sequence of ORF:</u>

ATGTTCGTGTTCCTGGTGCTCCTGCCTCTGGTGAGCAGCCAGTGCGTGAACCTGACC
ACCCGAACCCAGCTCCCACCAGCCTACACCAACAGCTTTACACGGGGCGTGTACTAC
CCTGACAAGGTGTTCAGATCTAGCGTCCTGCACAGCACTCAGGACCTCTTCCTGCCG
TTCTTCAGCAACGTGACATGGTTCCACGCCATCCACGTGAGCGGCACAAACGGAAC
CAAGCGGTTTGATAACCCCGTCCTGCCATTCAATGATGGAGTTTACTTCGCCAGTAC
CGAGAAGAGTAACATCATCCGGGGCTGGATCTTCGGCACCACCCTGGATAGCAAAA
CACAGAGCCTCCTGATCGTGAACAATGCCACGAACGTCGTGATCAAGGTGTGCGAG
TTCCAGTTTTGCAATGATCCTTTCCTGGGTGTGTACTACCACAAGAACAACAAGAGC
TGGATGGAAAGCGAGTTCAGAGTCTACAGCAGCGCCAACAACTGCACATTCGAGTA
CGTCTCTCAGCCTTTTCTGATGGACCTTGAGGGGAAACAAGGCAACTTCAAGAACCT
GAGAGAATTCGTGTTCAAGAACATCGACGGCTACTTCAAAATCTACTCCAAGCACAC
ACCCATCAACCTGGTCCGGGACCTCCCTCAGGGCTTCAGCGCCCTGGAACCCCTGGT
CGACCTGCCCATAGGCATCAACATAACGCGGTTCCAAACCCTGCTGGCCCTGCATAG
ATCCTACCTGACTCCTGGCGACAGCAGCAGCGGATGGACCGCCGGAGCTGCAGCCT
ACTATGTGGGCTACCTGCAACCTAGAACCTTCCTGCTGAAGTACAACGAGAACGGC
ACAATCACAGACGCCGTCGACTGCGCCCTGGACCCTCTCTCTGAGACAAAGTGCACC
CTGAAGTCCTTCACCGTGGAAAAGGGCATCTACCAGACCAGCAACTTCCGGGTGCA
GCCTACAGAGAGCATCGTGCGATTTCCAAACATTACCAACCTCTGCCCCTTCGGCGA
GGTGTTTAACGCCACAAGATTTGCCTCCGTTTACGCCTGGAATAGAAAGAGAATCAG
CAATTGTGTGGCCGACTACTCCGTGCTGTATAACAGCGCCTCTTTCAGCACCTTCAA
GTGCTACGGCGTTTCCCCAACAAAGCTGAATGACCTGTGCTTCACCAACGTGTACGC
CGACTCCTTCGTAATTAGAGGCGATGAGGTGCGGCAGATCGCACCAGGCCAGACCG
GTAAGATCGCTGACTACAACTATAAGCTGCCTGATGATTTTACAGGCTGCGTGATCG
CCTGGAACTCTAACAACCTGGATAGCAAGGTGGGCGGCAACTACAACTACCTGTAC
CGGCTGTTTCGCAAGTCTAACCTGAAACCTTTCGAGAGAGACATCTCCACAGAGATC
TACCAGGCCGGTTCTACACCTTGTAACGGGGTGGAAGGCTTCAACTGTTACTTCCCT
CTGCAAAGCTACGGCTTCCAGCCTACCAATGGAGTCGGCTACCAGCCATACCGGGT
GGTCGTGCTGTCCTTCGAGTTACTCCACGCCCCCGCCACCGTCTGCGGTCCTAAGAA
GTCCACCAATCTGGTTAAGAACAAATGCGTGAACTTCAACTTCAACGGCCTGACCGG
GACCGGCGTGCTGACCGAAAGCAACAAAAAGTTCCTCCCCTTCCAGCAGTTCGGCC
GTGATATCGCTGACACCACAGATGCCGTCAGAGATCCACAGACCCTGGAAATCCTG
GATATTACACCCTGCTCCTTCGGAGGAGTTTCTGTGATCACCCCCGGGACCAATACC
AGCAACCAGGTGGCTGTGCTGTACCAAGATGTTAACTGCACCGAGGTTCCTGTGGCC
ATCCACGCCGATCAGCTGACACCTACTTGGAGAGTGTACTCCACTGGCTCCAATGTG
TTCCAGACCAGGGCCGGATGTCTGATCGGCGCCGAGCACGTGAATAACAGTTACGA
GTGCGACATCCCTATCGGCGCCGGCATCTGTGCCAGCTACCAGACCCAGACAAACA
GCCCTGGGTCTGCTTCCTCTGTAGCTAGCCAGAGCATCATCGCCTACACCATGAGCC
TGGGCGCAGAGAACAGCGTGGCCTATTCCAACAACTCTATCGCCATTCCCACCAACT
TTACAATTAGCGTCACAACAGAGATCCTGCCCGTGAGCATGACCAAGACCAGCGTG
GACTGTACAATGTACATCTGTGGCGACAGCACTGAATGCAGCAACCTGCTGCTGCAA
TACGGCTCCTTTTGCACCCAACTGAACCGGGCGCTGACCGGAATCGCCGTGGAACA

GGACAAAAATACCCAGGAGGTGTTCGCCCAAGTGAAGCAGATCTACAAGACCCCAC
CTATCAAGGACTTCGGCGGCTTTAACTTTAGCCAGATTCTCCCTGATCCTTCTAAGCC
TAGCAAGCGGAGCTTTATCGAGGATCTGCTGTTCAACAAGGTCACCCTGGCCGATGC
CGGCTTTATCAAACAGTATGGCGATTGCCTGGGCGACATAGCCGCCAGAGATCTGAT
CTGCGCCCAGAAATTCAACGGCCTGACAGTTCTCCCACCTCTGCTGACCGACGAGAT
GATCGCTCAGTACACCTCTGCCCTGCTGGCTGGCACCATCACATCTGGGTGGACATT
TGGCGCCGGCGCCGCCCTGCAGATCCCCTTTGCCATGCAGATGGCCTATAGATTCAA
CGGAATCGGCGTGACCCAGAACGTGCTGTATGAAAACCAGAAGCTGATCGCTAACC
AGTTCAATTCTGCCATCGGCAAGATCCAGGACTCCCTCTCCTCTACCGCCAGCGCCC
TGGGCAAACTGCAGGACGTGGTGAATCAGAACGCCCAAGCCCTGAACACCCTGGTG
AAGCAGCTCAGCAGCAATTTTGGCGCCATCAGCTCTGTGCTGAACGATATCCTGTCT
AGACTGGACCCTCCAGAAGCCGAAGTCCAGATCGATAGACTGATCACAGGCAGACT
GCAGTCCCTGCAAACCTACGTGACCCAACAGCTGATCAGGGCCGCTGAAATAAGAG
CCAGCGCCAATCTCGCCGCTACCAAGATGTCCGAGTGTGTGCTGGGACAGTCTAAAC
GCGTTGACTTCTGCGGCAAAGGCTATCACCTGATGAGCTTCCCCCAGAGCGCGCCGC
ACGGCGTGGTGTTCCTGCATGTGACATACGTGCCTGCCCAAGAGAAGAATTTCACAA
CCGCCCCTGCCATCTGCCACGACGGCAAGGCCCACTTCCCAAGAGAGGGCGTTTTCG
TTTCCAATGGCACACACTGGTTCGTGACACAAAGAAACTTCTACGAACCCCAGATTA
TCACCACCGACAACACCTTCGTGAGTGGCAATTGTGACGTGGTCATCGGAATCGTGA
ACAACACAGTGTACGACCCTCTGCAACCTGAGCTGGACTCTTTTAAGGAAGAGCTGG
ACAAGTACTTTAAAAACCACACCAGCCCCGATGTGGACCTGGGCGACATCAGTGGC
ATTAACGCCAGCGTGGTGAACATCCAAAAGGAAATCGACAGACTGAACGAGGTGGC
CAAGAACCTGAACGAGTCCCTGATCGACCTGCAGGAGCTCGGCAAATACGAGCAGG
GATCCGGATACATCCCCGAGGCCCCCAGAGATGGCCAGGCCTACGTGCGGAAGGAC
GGCGAGTGGGTACTGCTGAGCACATTCCTGGGCAGATCCCTGGAGGTGCTGTTCCAG
GGCCCAGGCCATCACCACCATCACCACCATCATAGCGCCTGGTCCCACCCCCAGTTC
GAGAAGGGCGGCGGTAGTGGAGGGGGCGGATCTGGCGGCTCAGCTTGGAGCCACCC
CCAGTTCGAAAAGTGA

Citation: Wrapp, Daniel, et al. "Cryo-EM structure of the 2019-nCoV spike in the prefusion conformation." Science 367.6483 (2020): 1260-1263.

pCAGGs plasmid: https://www.addgene.org/vector-database/2042/
